# ALS-associated exitron splicing produces UBQLN2 isoforms with distinct properties

**DOI:** 10.64898/2026.09.08.750250

**Authors:** Adenine Si-Hui Koo, Jingjing Zhou, Natasha M. Méndez-Albelo, Yuchun Wei, Wei Xu, Xinyu Zhao, Colin Dewey, Randal S. Tibbetts

**Affiliations:** Department of Radiation Medicine, School of Medicine and Public Health, University of Wisconsin-Madison, Madison, Wisconsin, 53705, USA; Cellular and Molecular Biology Training Program, University of Wisconsin-Madison, Madison, Wisconsin 53705, USA; McArdle Laboratory for Cancer Research, University of Wisconsin-Madison, Madison, Wisconsin 53705, USA; Carbone Comprehensive Cancer Center, University of Wisconsin-Madison, Madison, Wisconsin 53705, USA; Waisman Center, University of Wisconsin-Madison, Madison, Wisconsin 53705, USA; Department of Neuroscience, School of Medicine and Public Health, University of Wisconsin-Madison, Madison, Wisconsin 53705, USA; Molecular and Cellular Pharmacology Training Program, University of Wisconsin-Madison, Madison, Wisconsin 53705, USA; Department of Radiology and Medical Physics, University of Wisconsin-Madison, Madison, Wisconsin 53705, USA; Department of Biostatistics and Medical Informatics, University of Wisconsin-Madison, Madison, Wisconsin 53705, USA

## Abstract

X-linked amyotrophic lateral sclerosis (ALS) implicated Ubiquilin 2 (UBQLN2) is expressed from a single exon. Here, we show that human UBQLN2 mRNA is alternatively spliced by virtue of a cryptic exonic intron (exitron) spanning the 5’ untranslated region and the 5’ end of *UBQLN2* coding sequence. Splice-out of this exitron generates a spliced UBQLN2 (UBQLN2-Sp) transcript that is translated from codon M243 to produce an N-terminally truncated UBQLN2 isoform (UBQLN2-M243) with reduced stability, diminished proteasome targeting, and altered aggregation behavior following the introduction of ALS mutations. The RNA-binding proteins SRSF1 and PTBP1 control *UBQLN2* splicing through binding to splice donor-proximal motifs, while TDP-43 was implicated as an indirect splicing repressor. *UBQLN2* splicing was elevated and inversely correlated with *UBQLN2* gene expression in the medial motor cortex of male ALS patients. These findings suggest that alternative splicing regulates *UBQLN2* gene dosage and function, which may impact UBQLN2-ALS proteinopathy.

**GRAPHICAL ABSTRACT:** Created in https://BioRender.com

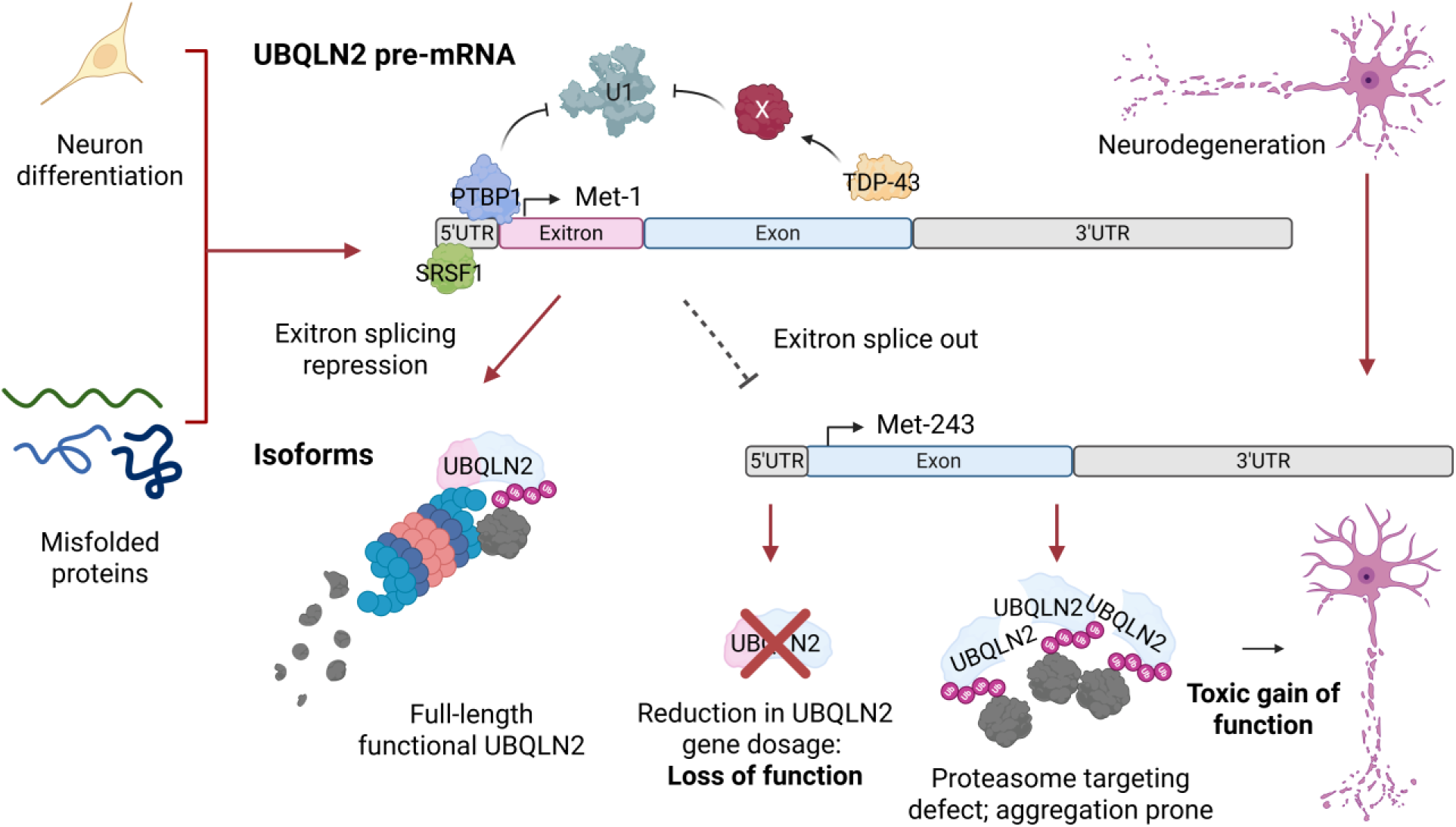

## INTRODUCTION

Amyotrophic lateral sclerosis (ALS) is a fatal neurodegenerative disease (ND) that affects upper and lower motor neurons with a wide range of clinical manifestations. Current treatment options are limited, and patients generally succumb to the disease within three to five years from symptom onset ^1^. Despite rigorous research, the causation and pathogenesis of ALS fail to be delineated. As reviewed in ^2^, ALS involves complex interplay between genetic components and environmental factors. Rare allelic variants from more than 40 genes have been identified as major risk factors, rendering ALS a polygenic disease which is further complicated by pleiotropic genes and incomplete penetrance (https://www.als.org/).

Approximately 5-10% of ALS patients have a known family history of the disease ^3–6^. Mutations in four genes account for ∼70% of these familial ALS cases ^2,3^: chromosome 9 open reading frame 72 (*C9ORF72*) ^7,8^; superoxide dismutase 1 (*SOD1*) ^9^, *TARDBP* which encodes for transactive response DNA/RNA binding protein of 43 kDa (TDP-43) ^10^, and fused in sarcoma (*FUS*) ^11,12^. Both TDP-43 and FUS are RNA binding proteins (RBPs) which are involved in RNA processing predominantly within cell nuclei as reviewed in ^13,14^. ALS-associated mutations in TDP-43 and FUS interfere with their nuclear localization and phase separation dynamics, leading to the formation of cytosolic aggregates and splicing dysregulation of their target genes ^4,11,12,15^. In fact, TDP-43 sequestration into cytoplasmic inclusions is observed in 97% of sporadic ALS and 45% of sporadic frontotemporal dementia (FTD) ^16,17^. As a member of the heterogeneous nuclear ribonucleoproteins (hnRNPs), TDP-43 acts as a crucial repressor of cryptic exon splice-in ^16^. TDP-43 loss of function either through null mutation or cytoplasmic retention leads to the inclusion of cryptic exons in mature mRNA transcripts which may contain premature termination codons (*UNC13A*-CE) ^18,19^ or alternative polyadenylation signals (*Stathmin-2*-Ex2a) ^20,21^ that lead to truncation of full-length protein coding sequences and loss of function of the target protein. FUS, on the other hand, associates with minor spliceosomes involved in the splicing of U12-dependent introns ^22^.

Splicing dysregulation has emerged as an important hallmark in ALS as well as other NDs ^15,23^. For instance, mis-splicing of *MAPT* gene is associated with tauopathy ^24–26^, while misplicing of *amyloid precursor protein* (*APP*) ^27,28^, *presenilin-1* and *presenilin-2* (*PSEN1* and *PSEN2*) ^29,30^, *Apolipoprotein E4* (*APOE4*) ^28^ and its receptor *APOE2R* ^31^ were found to either directly or indirectly contribute to Alzheimer’s disease (AD). In Parkinson’s patients, alternative *α-synuclein* (*SNCA*) splicing produced shorter translational products with altered aggregation characteristics compared to full-length α-synuclein ^32^. Finally, the incomplete splicing of *huntingtin* (*HTT*) Exon 1, which encodes the pathogenic CAG repeats, leads to the production of the highly toxic Exon 1-polyglutamine peptide in Huntington’s Disease (HD) patients ^33^. Owing to these causal links between splicing dysregulation and NDs, RBPs have come into the spotlight as potential diagnostic and therapeutic targets.

Repressor and enhancer RBPs play an important role in establishing and maintaining a steady pool of mature mRNAs specific for each cell lineage and in different stages of development ^34,35^. Polypyrimidine tract binding protein 1 (PTBP1) – also a member of the hnRNP family prevalently known for its repressor role in alternative splicing – is a ubiquitously expressed splicing regulator that binds to polypyrimidine-rich regions, either exonic or intronic, to influence splice site selection through steric blockage of spliceosome assembly or RNA looping ^36–40^. In the developing brain, downregulation of PTBP1 and corresponding upregulation of neural PTBP2 accounts for ∼25% of neuron-specific alternative splicing and critically contributes to neural differentiation and synaptogenesis ^35,41–44^. The importance of PTBPs in NDs is exemplified by PTBP1 downregulation in Parkinson’s disease ^45^ and its upregulation in AD, which is associated with enhanced Exons 7 and 8 inclusion in *APP* mRNA ^46^. On the other hand, sequestration of serine and arginine-rich splicing factor 1 (SRSF1), on Intron-1 retained C9ORF72 transcripts, was linked to enhanced nuclear export ^47,48^. SRSF1 depletion prevented the cytoplasmic translation of C9ORF72 repeats into dipeptide-repeat proteins, which in turn conferred neuroprotection ^47,48^. A more recent study has demonstrated that SRSF1 plays a role in the selection of a downstream splice donor (SD) of C9ORF72-Intron 1 to exonize the expanded repeats and promote the nuclear export of the repeat-containing transcripts ^49^. Nonetheless, SRSF1’s role in NDs is emerging.

*Ubiquilin 2* (*UBQLN2*) is a monoexonic gene arising from retroposition of UBQLN1 reverse-transcribed mRNA into the X chromosome of eutherians ^50^. UBQLN2 – a 624 aa member of the ubiquilin family of Ub chaperone proteins – is unique to placental mammals. Ubiquilins are organizationally defined by the presence of N-terminal ubiquitin-like (UBL), and a C-terminal ubiquitin-associated (UBA) domain flanking an array of centrally located, low complexity, STI1-like repeats that mediate liquid-liquid phase separation (LLPS) and binding to hydrophobic client proteins ^51–53^. A generic model for ubiquilin function holds that they engage polyubiquitylated substrates through their UBA domain and deliver them for degradation via UBL-dependent interactions with the 26S proteasome in the ubiquitin-proteasome system (UPS) ^54–56^. *UBQLN2* and its related paralogs, *UBQLN1* and *UBQLN4* are thought to function semi redundantly in mislocalized membrane protein triage, autophagy, and protein degradation (reviewed by ^57^), however, neurological and proteostasis phenotypes seen in *Ubqln2* knockout mice and UBQLN2-deficient cells indicate that it executes non-redundant functions as well ^58^.

Missense mutations clustered in a proline-rich repeat (PRR) that is unique to UBQLN2 cause dominant X-linked ALS, FTD, and/or spastic paraplegia ^59–63^. ALS-associated mutations in *UBQLN2* promote its aggregation ^64^ and have been linked to UPS dysfunction through impaired HSP70 binding ^54^, stress granule (SG) formation anomalies ^53,65^, autophagy defects ^66–70^, endoplasmic reticulum-associated protein degradation (ERAD) defects ^71–73^, lipid droplet dysregulation ^74^, reduced NF-kB activation ^75^, axon guidance anomalies ^76–78^, and other defects as reviewed in Ref. ^79,80^. Both pathogenic UBQLN2 and TDP-43 inclusions are rarely coincident in post-mortem ALS/FTD-UBQLN2 patient brain and spinal cord ^81,82^. However, WT UBQLN2 co-aggregated with poly-GA (C9ORF72 dipeptide-repeat) with low phosphorylated TDP-43 and p62 signals in human hippocampus, whereas UBQLN2-ALS mutants are more prone to co-aggregation with phosphorylated TDP-43 and p62 ^81^. Interestingly, clinical disease is more strongly correlated to the pattern of TDP-43 inclusion pathology than UBQLN2 pathology, suggesting that UBQLN2 macroinclusions are insufficient for disease manifestation ^82^.

Here, we show that human *UBQLN2* (*HsUBQLN2*) contains an unannotated exitron – a protein coding intron – whose removal generates N-terminally-truncated isoforms which alter the stability, aggregation propensity, and proteosome association of full-length UBQLN2. *UBQLN2* exitron splicing is primate-specific and regulated by *cis*-acting splicing elements which are bound by *trans*-acting RBPs, including but not defined solely by the splice sites and a series of polypyrimidine (Py) tracts in the *UBQLN2* 5’ untranslated region (5’UTR). Our finding suggests that regulated exitron removal has the potential to modulate *UBQLN2* gene dosage and function, with potential impacts on UBQLN2-mediated proteostasis, as demonstrated by higher incidence of exitron splicing in ALS patient transcriptomes.

## MATERIALS AND METHODS

### RNA sequencing analysis and alignment

PacBio-UHRR (Universal Human Reference RNA) dataset was accessed from Pacific Biosciences Datasets at https://www.pacb.com/connect/datasets. ONT-SGNex-HepG2 data set was accessed from The Singapore Nanopore Expression Data Set at https://registry.opendata.aws/sgnex. The bam files of interest were viewed on UCSC genome browser against UCSC hg38 reference. Illumina-TargetALS dataset was obtained from the Target ALS Foundation as fastq files (Sample Source = Cortex_Motor_Medial; C9orf72 Repeat Expansion (Data from CUMC) = No/Unknown). The splice junctions, SA1 and SA2, detected from the long-read data were annotated manually and appended to the GENCODE v48 genome annotation file. The reads were aligned to UCSC hg38 reference genome (https://hgdownload.soe.ucsc.edu/goldenPath/hg38/bigZips/hg38.fa.gz) by the splice aware aligner – STAR (Spliced Transcripts Alignment to a Reference) ^83^. STAR-aligned bam files (--quantMode TranscriptomeSAM) were used as inputs for *UBQLN2* spliced isoform expression estimation by RSEM (RNA-Seq by Expectation Maximization) algorithm ^84^. Because RNA degradation and 3’ read bias can confound the analysis of splicing events occurring at the 5’ end of the *UBQLN2* transcript, only sequencing samples with an RNA Integrity Number (RIN) ≥ 5.5 were included in downstream analysis. For samples with missing RIN, its RIN was imputed from a linear regression model which was trained on the samples with known RIN using the read start position distribution (RSPD) and fragment length statistics estimated by RSEM. The model was evaluated using leave-one-out cross-validation (LOOCV) which resulted in an r^2^ of 0.64 for predicted vs. observed RIN values. See Sup. Table 1 for the ExternalSampleID of the samples which passed the filtering criteria. UBQLN2-Spliced isoform expression was quantified by the sum of the estimated RSEM Isoform Percentage (IsoPct) of SA1 and SA2 transcripts. The samples were grouped by Sex (Female vs Male) and Subject/Clinical Group (ALS Spectrum MND vs Non-Neurological Control). Wilcoxon tests were performed which agreed with the results from linear modeling of logit transformed spliced isoform expression values. *UBQLN2* gene expression was quantified as transcripts per million (TPM) from RSEM and a linear regression analysis was performed on the relationship between *UBQLN2* TPM and spliced isoform expression.

For sequence alignment, reference mRNA sequences of *UBQLN2* from *Homo sapiens* (NM_013444.4), *Mus musculus* (NM_018798.2), *Pan troglodytes* (XM_001148687.7), *Pongo abelii* (XM_002831718.6), and *Pongo pygmaeus* (XM_054472304.2) were downloaded from NCBI on 15-Sep-2025. Sequences between the 5’ untranslated region (5’ UTR) and the translational start site (TSS, Met-1) were aligned manually. The UBQLN2 mRNA sequences at the selected region were identical for all primates and hence not shown in the alignment figure to omit redundancy.

### Cell culture and treatment

HeLa, HEK293T, and U-2 OS cell lines were obtained from the American Type Culture Collection (ATCC). HeLa and HEK293T cells were grown in DMEM medium (Corning, 10-013-CV) whereas U-2 OS were grown in McCoy’s 5A medium (Corning, 10-050-CV). Murine 4T1 (BALB/cfC3H) and MC38 (C57BL/6) cells were originally obtained from ATCC. 4T1 was maintained in DMEM whereas MC38 was maintained in RPMI medium. All growing medium was supplemented with 10% fetal bovine serum (GeminiBio 900-108-500) and 1% Penicillin/Streptomycin (Corning, 30-002-CI). The cells were incubated at 37°C in 5% CO_2_ unless otherwise stated.

For heat shock exposure, HeLa cells were placed in 42°C incubator supplemented with 5% CO_2_ for two hours followed by recovery at 37°C before being harvested. Where indicated, 50 µM MG132 or 100 nM bafilomycin A1 (reconstituted in DMSO), was added to transfected HeLa cells or CRISPR-generated UBQLN2-mNeon clones. Cells were harvested for Western blot analysis in modified RIPA buffer 4 h post-treatment.

### Lentiviral shRNA knockdown

Lentiviral vectors targeting the following genes were purchased from Sigma: UBQLN2 (cat# TRCN0000004377); SRSF1 (cat# TRCN0000001095), PTBP1 (cat# TRCN0000231420). The lentiviral TDP-43 plasmid was homemade by the following primers: TDP-43-pLKO.1-F: 5’- CCGGGCTCTAATTCTGGTGCAGCAACTCGAGTTGCTGCACCAGAATTAGAGCTTTTTG-3’ and TDP-43-pLKO.1-R: 5’- AATTCAAAAAGCTCTAATTCTGGTGCAGCAACTCGAGTTGCTGCACCAGAATTAGAGC-3’). The lentiviral FUS-3’UTR and FUS-CDS shRNA vectors were prepared as previously described ^85^. Non-targeting (shNT) vector (Addgene Plasmid #1864) was included as negative control. Lentiviral particles were produced by transient transfection of HEK293T cells with shRNA vectors, psPAX2 (Addgene plasmid #12260) and pCMV-VSV-G (Addgene plasmid #8454) in a ratio of 4:3:3 by jetPRIME^®^ as described ^85,86^. Lentiviral-infiltrating media harvested at 24 h and 52 h post-transfection were combined and centrifuged at 3,000 rpm for 15 min at 4°C. Viral supernatant was aliquoted 1 mL per tube before freezing at −80°C. HeLa cells were plated in 6-wells plate and transduced with 1 mL lentiviral supernatant per well at 70-80% confluency for 24 h, followed by selection with 2 µg/ml puromycin for 48-72 h. Cells were harvested in TRIzol reagent for splicing assays or modified RIPA buffer for Western blotting of proteins of interest.

### RNA extraction, cDNA syntheses, splicing assay, and Sanger sequencing

Total RNA was extracted from human cell lines (HeLa, HEK293T, and U-2 OS), mouse 4T1 and MC38 cells, as well as C57BL/6J mouse brain tissues by TRIzol reagent (Invitrogen 15596018) and reverse transcribed into cDNA using iScript™ gDNA Clear cDNA Synthesis Kit (Bio-Rad, 1725035) according to manufacturer’s protocols. RT-PCR was performed with GoTaq Green Master Mix (Promega M7123) with primers designed to amplify both full-length (FL) and spliced (Sp) UBQLN2 transcripts in human (Hs) or mouse (Mm). HsUBQLN2-F: 5’-AACCGCAGTCTTCATCACAG-3’; HsUBQLN2-Nter-R: 5’-CATGGATGCCATGCTGGATCAAGG-3’; HsUBQLN2-Sp-R2: 5’-

ATTGGGTAGTGGATCGCGAT-3’; MmUBQLN2-F: 5’-

GGAGGAGCCGCAGCCTTCAACTCTG-3’; MmUBQLN2-Nter-R: 5’-

CGGACGGTTCTGGCTTTTGATAAC-3’; MmUBQLN2-Sp-R2: 5’-

CGCCGCAGAGCATTGTAG-3’. Splicing primers amplified both UBQLN2-FL and UBQLN2-Sp amplicons which differ by 763 nt (Hs) and 781 nt (Mm) that were resolved by 2% w/v agarose gel electrophoresis. Gel images were acquired by Odyssey Fc/XF (LI-COR Biosciences). Five percent v/v DMSO was included to amplify *MmUBQLN2* splice products owing to high GC content.

Quantitative RT-PCR (qRT-PCR) of human samples was performed using HsUBQLN2-F and the following reverse primers: (HsUBQLN2-Nter-R4: 5’- AGAGCGCGAGGGAAGGAGGAAGG-3’; HsUBQLN2-Sp-R5: 5’- CTGGCAATTTCGAGTGTCTGCCTC-3’) with iTaq Universal SYBR® Green Supermix (Bio-Rad, 1725124) according to manufacturer’s protocols. GAPDH was included as loading control (GAPDH-F: 5’-AATCCCATCACCATCTTCCA-3’; GAPDH-R: 5’- TGGACTCCACGACGTACTCA-3’). For mouse samples, qRT-PCR was performed using MmUBQLN2-F and the following reverse primers (MmUBQLN2-Nter-R5: 5’- AAGAGGGCGCGAGGGAAGGAGGT-3’; MmUBQLN2-Sp-R6: 5’- CAGCGTCTGCCTCATTATATCTGG-3’). GAPDH was included as loading control (MmGAPDH-F: 5’-ACCTGCCAAGTATGATGA-3’; GAPDH-R: 5’- GGAGTTGCTGTTGAAGTC-3’).

For RT-PCR analysis of human and mouse UBQLN2 splicing reporters, the forward primer was located within the multiple cloning site of the peGFP-N1 backbone to specifically amplify transcripts from the reporters (peGFPN1-MCS-F: 5’- TCAGATCTCGAGCTCAAGCTTCG-3’). For qRT-PCR splicing assay involving mutated HsUBQLN2 reporter, a new set of reverse primers was designed to avoid the polypyrimidine tracts (HsUBQLN2-Nter-R6: 5’-AAGGAGGCACCGCCGCAGCG-3’; and HsUBQLN2-SA2-R8: 5’-CTGGGTTGTTGAGCAGGTGACTGA-3’). Additional reverse primer targeting region between SA1 and SA2 was included: HsUBQLN2-SA1-R9: 5’- GATTAGCCATAATGAGCTGCCTCA-3’.

Sanger sequencing was performed using BigDye Terminator v3.1 Cycle Sequencing Kit (Obtained from UWBC) with the following reverse sequencing primer 5’- CTAGATTGCTAAGAGCCAGGTC-3’ to verify the sequence identity of the amplicons. The PCR libraries were purified by using Axygen® AxyPrep MAG DyeClean Up Kit (Axygen MAG-DYECL-5) following manufacturer’s protocol. The sequencing products were resolved by capillary electrophoresis for reads by UWBC. The ab1 files were viewed in SnapGene Viewer.

### RNA immunoprecipitation (RNA-IP)

RNA-IP was performed essentially as described ^87^. Approximately 50 million cells were lysed in 1 mL ice-cold NET-2 buffer (50 mM Tris-HCl (pH 7.5), 150 mM NaCl, 0.05% v/v NP40) supplemented with 2 mM 1,4-dithiothreitol, 0.2 U/µL RNasin Plus (Promega, N2611), 20 mM sodium fluoride, 20 mM β-glycerophosphate and 2X protease inhibitor cocktail (Sigma, P8340-5ml; Thermo Scientific, 78438). The lysate was sonicated in five pulses of 3 s ON, 30 s OFF followed by three pulses of 10 s ON and 30 s OFF at an amplitude of 30% (Fisher Scientific, FB120). The lysate was cleared by centrifugation at 14,000 *x* g for 10 min, 4°C. The supernatant was incubated with 5 µg of either the targeted antibodies (PTBP1: ThermoFisher 32-4800; SRSF1-M: Santa Cruz sc33652; TDP-43: Proteintech 10782-2-AP) or normal mouse or rabbit IgG controls (Millipore 12-371; 12-370) for 1 h on a nutator mixer at 4°C before Protein A/G PLUS-Agarose bead suspension (Santa Cruz, sc-2003) was added (20 µl/µg of antibody) for overnight incubation. The beads were washed with NET-2 buffer five times. The immunoprecipitated RNA was extracted by TRIzol reagent and reverse transcribed into cDNA by iScript™ gDNA Clear cDNA Synthesis Kit. Log_2_ fold enrichment of UBQLN2 mRNA in the target RNA-IP sample versus the control IgG sample was quantified by qRT-PCR assay with primers flanking the 5’ splice site (HsUBQLN2-F and HsUBQLN2-Nter-R4). For TDP-43 RNA-IP, log_2_ fold enrichment of *STMN2* was included as a positive control: STMN2-Ex2a-F: 5’- TTTGGCTCTCTGTGTGAGCA-3’; STMN2-Ex2a-R: 5’-CACAAGCCGCATTCACATTCA- 3’).

### iPSC-ExN differentiation and transcriptomic analysis

Human iPSCs were derived from healthy control male donors as in previous publications: GM00498-4 ^88^ and WC032i-6007-1 ^89^. Human iPSCs were maintained and the differentiation process was carried out according to the established protocols in ^90^. Human pluripotent stem cells (hPSCs) were thawed onto MEF feeder plates in hESC media supplemented with 10 µM ROCK inhibitor (Y-27632 dihydrochloride, Tocris #1254), passaged weekly onto new MEF plates by using 1 mg/ml collagenase Type IV (Thermo Fisher Scientific #17104019) in DMEM/F12. Human iPSCs were fed every day with hESC media (DMEM/F12 (Thermo Fisher Scientific, 11330032), KnockOut™ Serum Replacement (KOSR) (Thermo Fisher Scientific, 10828028), L-glutamine (Thermo Fisher Scientific, 25030081) and 2x FGF2 (Waisman Biomanufacturing, 8 ng/mL).

Neural differentiation of hPSCs was carried out using previously established dual SMAD inhibition-based protocol ^88–90^. hPSCs neural differentiation was induced 5 days after passaging of hPSCs onto MEFs by switching hESC medium to neural induction medium (NIM), consisting of 1:1 mixture of DMEM/F12:Neurobasal (Thermo Fisher Scientific 21103049), 1X N2 (Waisman Center), 1X L-glutamine, 1X Antibiotic-Antimyotic (Anti-Anti) (Thermo Fisher Scientific 15240062), 10 µM SB432542 (Biogems #3014193), 100 nM LDN193189 (Selleck Chemicals #S7507), and 5 µM XAV-939 (Selleck #3748). Cells were cultured in NIM for 9 days with a daily medium change with freshly added small molecules.

Cells were dissociated with TrypLE Express (Thermo Fisher Scientific #12605010) and replated in a 1:1 ratio onto Cultrex (R&D Systems) -coated plates in neural progenitor cell (NPC) medium (Neurobasal medium, 1X L-glutamine, 1X N2, 0.5X B27 without vitamin A (Thermo Fisher Scientific), 1X Anti-Anti), supplemented with 10 µM ROCK inhibitor. Cells were maintained for 6 days with daily change of NPC medium to transition from neuroepithelia to NPCs and reorganize into neural rosettes. At this stage, the NPCs could be dissociated with Accutase (Thermo Fisher Scientific), frozen in NPC freeze medium (90% FBS and 10% DMSO), and stored in liquid nitrogen.

The NPCs were thawed onto Cultrex-coated plates in the NPC media containing 10 µM ROCK inhibitor. For differentiation into excitatory neurons (ExNs), NPCs were plated at low cell density in the NPC medium supplemented with 10 µM ROCK inhibitor and 0.2 µM γ-Secretase Inhibitor XXI, Compound E (Calbiochem). On the next day, the media was changed into neuronal differentiation medium (NDM) (Neurobasal medium, 1X GlutaMAX (Thermo Fisher Scientific), 1X Anti-Anti, 1X N2, 0.5X B27 without vitamin A, 0.3% glucose, 20 ng/ml BDNF (Peprotech), 20 ng/ml GDNF (Peprotech), 500 ng/ml cAMP (Millipore-Sigma), and 200 µM ascorbic acid (Millipore-Sigma), supplemented with 0.2 µM Compound E. Post-mitotic neurons were maintained in NDM, with 0.5 vol of the medium change performed every 3 days. For RNA isolation and RT-qPCR analysis, NPCs were collected at 1 day after thawing and neurons were collected at 1 week or 4 weeks of differentiation. Cells were collected in TRIzol reagent (Thermo Fisher Scientific), then RNA was extracted using Direct-zol RNA Microprep Plus kit (ZYMON RESEARCH #R2062) following the manufacturer’s instructions. First strand cDNAs were synthesized with both oligo dT and random primers using PrimeScript™ RT Reagent Kit (Takara #RR037A), following manufacturer’s protocols. qRT-PCR was performed as described earlier with 15 ng cDNA template per reaction.

### Lipofectamine-mediated transfection

HeLa or HEK293T cells were seeded into 6-well plates, 38 mm glass bottom dishes, or 6 cm plates and transfected at 60-80% confluency using 3 µg (6-well plate, 38 mm glass bottom dishes) or 7.5 µg (6-cm plate) of plasmid DNA and the Lipofectamine 3000 reagent kit (Invitrogen L3000015) using the manufacturer’s recommendations. The culture medium was changed prior to the addition of transfection mixture, and the cells were incubated for 24 h for transgene expression. Cells were then harvested for splicing assays, Western blot analysis, immunofluorescence or live-cell imaging.

SRSF1 and PTBP1 overexpression was performed by co-transfecting the corresponding plasmids (pcDNA3.1-FLAG-SF2, Addgene Plasmid #99021; Myc-PTBP1, Addgene Plasmid #23024) with *HsUBQLN2* splicing reporter in a 5:1 mass ratio. Western blot and *RIF1* splicing assay ^87^ were used to assess the efficiency of overexpression.

### Cloning and QuikChange mutagenesis

The coding sequence of full-length UBQLN2 (UBQLN2-M1) and two of its isoforms (UBQLN2-M243 and UBQLN2-M446) were amplified from HEK293T cells using attB-flanked primers (UBQLN2-M1-F: 5’- GGGGACAAGTTTGTACAAAAAAGCAGGCTTAatggctgagaatggcgag-3’; UBQLN2-

M243-F: 5’- GGGGACAAGTTTGTACAAAAAAGCAGGCTTAatgatgcaagagatgatgag-3’; UBQLN2-M446-F: 5’- GGGGACAAGTTTGTACAAAAAAGCAGGCTTAatgtcaaacccaagagcaatg-3’; and UBQLN2-CDS-R: 5’- GGGGACCACTTTGTACAAGAAAGCTGGGTTttacgatggctgggagcc-3’) by using Phusion High-Fidelity PCR Master Mix with HF Buffer (Thermo Scientific F530L). The attB-PCR products were purified after being resolved on 1% w/v agarose gel by Zymoclean™ Gel DNA Recovery Kit (Zymo Research Corporation D4007) following manufacturer’s protocol. The purified attB-PCR products were subsequently cloned into pcDNA5-eGFP-FRT/TO plasmid vector (Addgene plasmid #19444) by Gateway recombination (Invitrogen 11789020 and 11791020) following manufacturer’s one-tube protocol with overnight 25°C incubation in thermocycler for both BP and LR reactions. After Proteinase K digestion, the reaction mixtures were transformed into DH5α competent cells (*ccdB* sensitive) and plated. After overnight incubation at 37°C, colonies were selected and purified plasmids were sequenced in entirety.

For splicing reporters, human and mouse *UBQLN2* gene sequences were amplified from HEK293T and C57BL/6J mouse genomic DNA respectively with an EcoRI-flanked forward primer and a BamH1-flanked reverse primer (HsReporter-F: 5’- GCGTGACGAATTCcagagttgctgggagtgcgcgc-3’; HsReporter-R: 5’-

CGATCGATGGATCCgatggctgggagcccagcagcctt-3’; MmReporter-F: 5’-

GCGTGAC<u>GAATTC</u>ggagacggcctgcaggacctgc-3’; MmReporter-R: 5’- CGATCGAT<u>GGATCC</u>gatggctgagagcccagcagcctc-3’). The amplified gene fragments were purified as described above and eluted in 16 µL of nuclease-free water. The purified fragments and peGFP-N1 vector (Clontech 6085-1) were subjected to *Eco*R1-HF (NEB R3101S) and *Bam*H1-HF (NEB R3136S) digestion for 1 h at 37°C in 10X rCutSmart Buffer (B6004S). The fragments of the correct size were excised, gel purified and ligated by T4 ligase (NEB M0202) in a molar ratio of 1:3 vector to insert (100 ng of vector to 140 ng of insert as calculated by NEBioCalculator) in a 10 µL ligation reaction. The ligation reactions were incubated at 16°C overnight, transformed into DH5α competent cells and plated. After overnight incubation at 37°C, colonies were selected and purified plasmids were sequenced in entirety.

To swap the 5’UTR of *HsUBQLN2* and *MmUBQLN2* splicing reporters, one microgram of each of the reporter plasmids was subjected to digestion with *Not*I (NEB R3189S), which cleaved at a conserved GCGGCCGC motif immediately upstream of the start codon and immediately downstream of the GFP cassette. The resulting HsUBQLN2^CDS^-GFP and MmUBQLN2^CDS^-GFP fragments and their corresponding vector-5’UTR fragments were resolved on 1% agarose gel, excised, and purified. We then ligated the heterologous UBQLN2^CDS^-GFP (insert) and vector-5’UTR-containing fragments in a molar ratio of 1:2 vector to insert (100 ng of vector to 125 ng of insert as calculated by NEBioCalculator) in a 20 µL ligation reaction. The ligated product was resolved on a 1% agarose gel. Fragments of the correct size were excised and purified before being transformed into DH5α competent cells and plated. Colonies were selected after overnight incubation at 37°C for plasmid purification. Purified plasmids were sequenced in entirety to screen for the correct insert orientation. The MmUTR-HsCDS reporter is 6800 bp in size, encoding a 57 nt shorter *HsUBQLN2* transcript than WT. The HsUTR-MmCDS reporter is 6899 bp in size, encoding a 57 nt longer *MmUBQLN2* transcript than WT, due to the difference in 5’UTR length.

For QuikChange Mutagenesis, primers containing the desired mutations (Sup. Table 2) were employed to amplify the target sequence using Phusion™ Hot Start II DNA Polymerases (2 U/µL) (Thermo Scientific F549L) following manufacturer’s recommendations. For primers targeting regions with high GC content (with melting temperature ≥ 65°C, 4% v/v DMSO was added into the reaction. Template plasmids were removed from the reaction mixture by Dpn1 digestion (Thermo Scientific ER1702) at 37°C for 4 h. *Dpn*1-digested mixture was transformed into DH5α or Stbl3 competent cells and plated for colony selection. All purified plasmids were sequenced by University of Wisconsin-Madison Biotechnology Center (UWBC) using the Oxford Nanopore platform. Guppy basecaller with the Super accurate (SUP) model was used for basecalling. pLannotate map which was provided by UWBC was subsequently screened for the desired mutations in SnapGene Viewer (version 6.0.2) for clone selection and plasmid amplification.

### Protein extraction and Western blotting

Cells were washed with 1X PBS and lysed on plate in 200 µL modified RIPA buffer (50 mM Tris-HCl (pH 7.5), 150 mM NaCl, 1 mM EDTA (pH 8), 1% sodium deoxycholate, 0.1% SDS, and 1% v/v Triton X-100) supplemented with 1X Halt™ Protease and Phosphatase Inhibitor Cocktail (Thermo Scientific 78446). The lysate was incubated on ice for 10 min, collected in 1.5 mL tubes, and sonicated in 5 pulses of 3 s ON, 5 s OFF at an amplitude of 30% (Fisher Scientific, FB120). The lysate was then mixed with 4X SDS sample buffer (200 mM Tris-HCl (pH 6.8), 40% glycerol, 8% SDS, 0.5% bromophenol blue and 10% beta-mercaptoethanol) at a ratio of 1:3. The samples were heated at 95°C for 5 min prior to freezing at −20°C for storage or loading directly for Western blotting.

For Western blotting, samples were separated by 12% SDS-polyacrylamide gel (SDS-PAGE). Proteins were transferred to 0.45 µm Immobilon®-FL PVDF Membrane (Millipore, IPFL00010) in Tris-glycine transfer buffer. The membranes were blocked with blocking solution (5% w/v milk in Tris-buffered saline, 0.1% v/v Tween 20 (TBST)) for 1 h before blotting with target primary antibodies overnight at 4°C. For UBQLN2-HPA antibody (Sigma HPA006431), the blocking reagent was replaced with 1% w/v bovine serum albumin (BSA) in TBST or 1X RapidBlock™ solution (VWR 97064-124). The source and the dilution of the primary antibodies used were listed as followed: UBQLN2-CST (Cell Signaling Technology 85509S, 1:1000); SRSF1 (Santa Cruz sc33652, 1:100); alpha-tubulin (Sigma-Aldrich T6199, 1:1000); vinculin (Santa Cruz sc73614, 1:1000); MCM2 (Abcam ab4461, 1:2000); PTBP1 (ThermoFisher 32-4800, 1:1000); TDP43 (Proteintech 10782-2-AP, 1:1000); GFP (Santa Cruz sc9996, 1:100); UBQLN2-HPA (Sigma HPA006431, 1:250); mNeonGreen-Tag (Cell Signaling Technology 55074, 1:1000), HA-Tag (Cell Signaling Technology 3724S, 1:1000). After primary antibody incubation, the membranes were washed 3 X 5 min with TBST and incubated with LI-COR IRDye secondary antibodies (IRDye 800CW goat anti-rabbit or IRDye 680RD goat anti-mouse) at a dilution of 1:10000 in blocking solution for 45 min at room temperature. Membranes were washed 3 X 5 min with TBST, and images were acquired using Odyssey XF (LI-COR Biosciences). The exported images were then analyzed and organized with ImageStudio software (v5.2, LI-COR Biosciences).

### CRISPR-Cas9 nucleofection

Alt-R™ CRISPR-Cas9 crRNA (Hs.HC9.FFFH5766.AA) and Alt-R™ CRISPR-Cas9 tracrRNA (IDT 1077024) were resuspended in IDTE buffer to a final concentration of 200 µM. Alt-R™ HDR Donor Block (CD.HC9.SJTY5528+) was resuspended in IDTE buffer to a final concentration of 500 ng/µL according to IDT recommendations. Equimolar concentrations of crRNA and tracrRNA were mixed in a sterile tube to a final duplex concentration of 100 µM. The crRNA:tracrRNA duplex mixture was incubated at 95°C for 5 min and cooled down to room temperature on benchtop. The ribonucleoprotein (RNP) complex was prepared by mixing 120 pmol of crRNA:tracrRNA duplex and 104 pmol of Alt-R™ S.p. HiFi Cas9 Nuclease V3 (IDT 1081060) in 1X PBS to a total volume of 5 µL, followed by 20 min incubation at room temperature.

Amaxa™ 4D-Nucleofector™ protocol for MCF7 for 4D-Nucleofector™ in suspension was used. Ten million HeLa cells were harvested by trypsinization. 4D-Nucleofector™ solution for the 20 µL Nucleocuvette™ Strip was prepared according to manufacturer’s protocol by mixing 16.4 µL of Nucleofector™ Solution and 3.6 µL of supplement. HeLa cell pellet was resuspended carefully at room temperature with the 4D-Nucleofector™ solution. RNP complex and 1 µg of the Alt-R™ HDR Donor Block was added into the cell suspension before the whole mixture was transferred into the 20 µL Nucleocuvette™ Strip. Nucleofection was performed on LONZA 4D-Nucleofector by using the default specification for HeLa cells. After nucleofection, 75 µL of pre-warmed complete DMEM medium was used to gently resuspend the cells. Resuspended cells were added to a well on 24-wells plate containing 1 mL pre-warmed medium which was pre-equilibrated in a humidified 37°C/5% CO_2_ incubator. The cells were allowed for recovery for 2 days before being transferred to a larger vessel. At 5-7 days after nucleofection, mNeon positive cells were single-cell sorted into 96-wells plate by BD FACSDiscover S8 "Sparky". The cells were expanded to 6-wells before being harvested. Genomic DNA from the clones were extracted in QuickExtract DNA Extraction Solution (Biosearch Technologies) according to manufacturer’s protocol. Genotyping of the clones were performed with the following primers whose relative positions were shown in Sup. Fig. 3A. HsUBQLN2-Cter-F2: 5’- CAGCAACAACTGGAACAGCTC-3’; HsUBQLN2-Cter-R2: 5’- GCATCATATTAGACTTACAACTAGAC-3’; mNeon-R: 5’- CCATCATTTGGATTGCCGGTGC-3’. Insertion of mNeon upshifted the amplicon from HsUBQLN2-Cter-R2 by 753 nt which was resolved by 2% w/v agarose gel electrophoresis. Gel images were acquired by Odyssey Fc/XF (LI-COR Biosciences).

### Immunofluorescence staining, live-cell Imaging, aggregates quantification

Cover slips (Fisher Scientific 12-545-83) were placed in each well of a 12-well plate, rinsed once with 75% ethanol and twice with 1X phosphate-buffered saline (PBS). Cells were seeded onto cover glass in each well, fixed with 4% w/v paraformaldehyde (PFA) for 15 min, permeabilized with 0.2% v/v Triton X-100 for 10 min and then blocked with 3% w/v BSA for an hour at room temperature. The cover slips were then transferred to an improvised humidity chamber for primary antibodies incubation at 37°C for 1 h or at 4°C overnight. The source and the dilution factor of the antibodies used were indicated here: alpha-tubulin (Sigma-Aldrich T6199, 1:500); proteasome 19S S5A/ASF antibody (Abcam ab154935, 1:100). The cover glasses were washed 5 X 1 min with 0.05% v/v PBST (PBS, 0.05% v/v Tween-20) before incubating with Alexa Fluor™ secondary antibodies (Thermo Fisher #A11032, #A32733) at a dilution of 1:1,000 for 45 min at room temperature. The cover slips were washed with 0.05% PBST for 3 X 1 min followed by 1X PBS for 2 X 1 min before being mounted on glass slides with DAPI-containing mounting medium (Vector H-1200). Images were acquired by Nikon AX confocal microscopes under the desired objectives and organized using NIS-Elements Advanced Research. For live-cell imaging, the cells were plated into 38 mm glass bottom dishes and stained with 1 µg/mL of Hoechst 33342 for 15 min before being imaged in humidified CO_2_ chamber supplemented with 5% CO_2_ throughout the experiment.

For aggregate quantification, GFP channel of the confocal images were retrieved and converted into a mask in Fiji. Aggregates were segmented and quantified by Analyze Particles-Shape Descriptors. Aggregates smaller than 0.2 µm^2^ were excluded for roundness quantification as they give a constant roundness value. Roundness of the aggregates were estimated by the following equation:

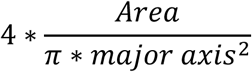

The results were tabulated by a custom script in MATLAB and plotted in Prism (version 10.6.1). For S5A colocalization, S5A signal mask was generated in Fiji. For each image, the Pearson correlation coefficient of S5A and its corresponding GFP mask was calculated by a custom script in MATLAB.

### Statistics

All statistical calculations were performed by Prism (version 10.6.1) following the recommended parameters. The sample sizes and tests performed were listed in the figure legends. The resulting *p*-values were printed on the figures.

## RESULTS

### Human *UBQLN2* contains an unannotated exitron

*Homo sapiens UBQLN2* (*HsUBQLN2*) is annotated as an intronless gene (hg38 coordinates chrX:56,563,627-56,567,868) that produces a single 624 amino acid (aa) translational product from the first AUG codon (UBQLN2-M1). We queried several long-read sequencing databases derived from two major long-read sequencing platforms (the Universal Human Reference RNA/UHRR sequenced on Pacific Biosciences (PacBio) Sequel II and Revio systems, as well as HepG2 cell RNA sequenced on Oxford Nanopore Technologies (ONT) MinION (FLO-MIN106D) ^91^. These analyses identified an exitron spanning the 5’ UTR and the coding sequence (CDS) of *HsUBQLN2*. We adopted the exitron nomenclature (exonic intron) which involves alternative splicing within a protein-coding exon through the activation of cryptic splice sites as originally described by Marquez *et al.* in 2015 ^92^. The putative *HsUBQLN2* exitron starts at a consensus SD sequence at genomic position chrX:56,563,741 and ends with two alternative SAs at chrX:56,564,449 (SA1) or chrX:56,564,503 (SA2) of the *HsUBQLN2* gene (Fig. 1A, annotated by *red* and *black arrows* respectively). Splicing out of this exitron results in the production of at least two 5’-truncated UBQLN2 transcript variants depending on SA usage: UBQLN2-SA1 (Fig. 1A, *red arrow*) and UBQLN2-SA2 (Fig. 1A, *black arrows*) that lack much of the 5’ UTR and the sequences encoding for either the first 193 (SA1) or 210 (SA2) amino acids of the UBQLN2 CDS. In both transcripts, the first available AUG codon is in-frame with the full-length UBQLN2 CDS. Any translation products produced from these transcripts lack the proteosome-targeting UBL domain and the first chaperone-binding STI1 domain, likely leading to altered functional properties relative to UBQLN2-M1 (Fig. 1A, protein domains). Screening of the short-read RNA-sequencing data from TargetALS RNA sequencing datasets based on Illumina sequencing system (HiSeq 2500) also detected reads that mapped across the splice junctions of both UBQLN2-SA1 and UBQLN2-SA2 transcripts (Fig. 1A, ExternalSampleID: CGND-HRA-00013; ExternalSubjectID: NEUEL133AK6), suggesting that the identified variants were not artefacts resulting from a particular sequencing approach/platform.

**Fig. 1.**
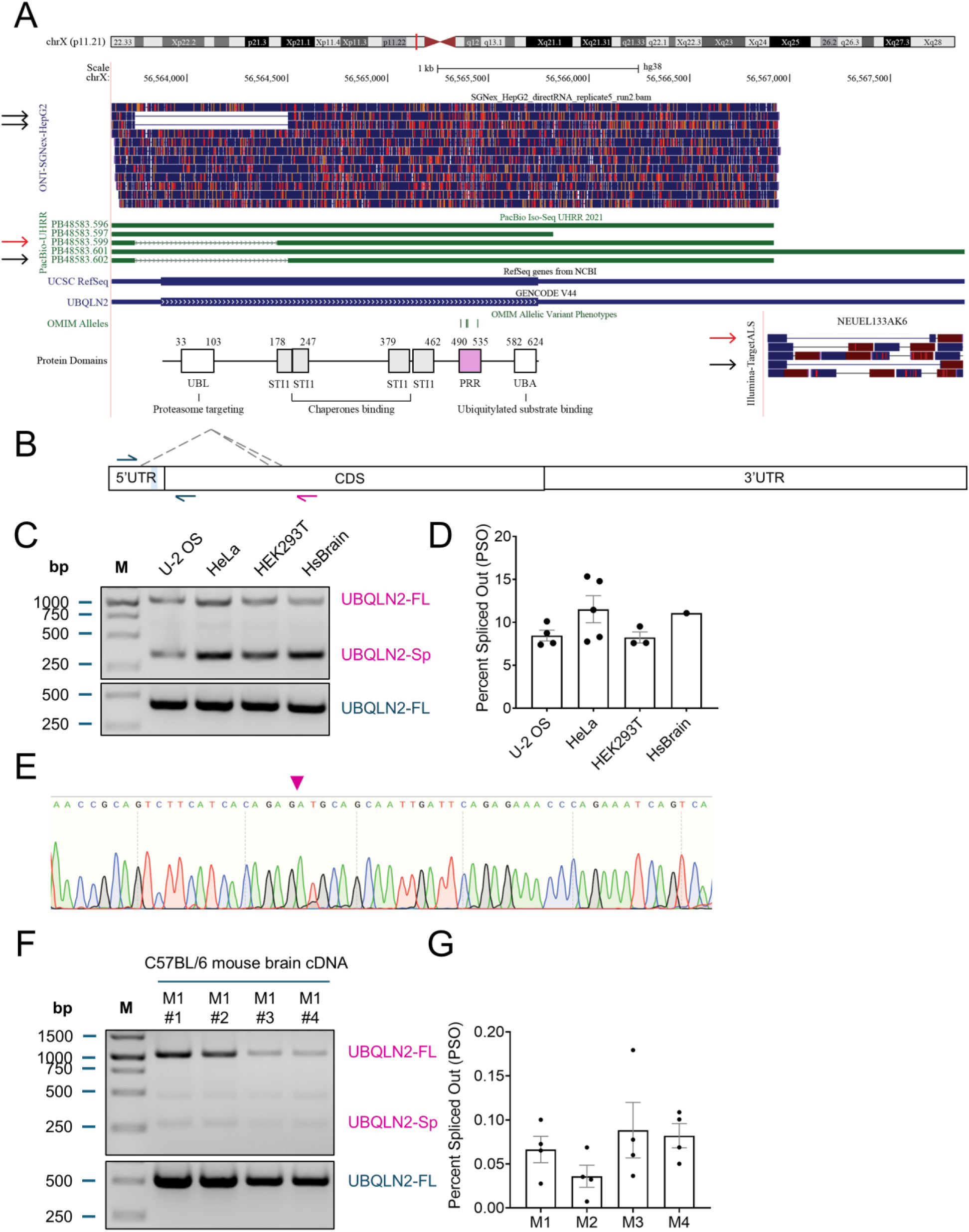
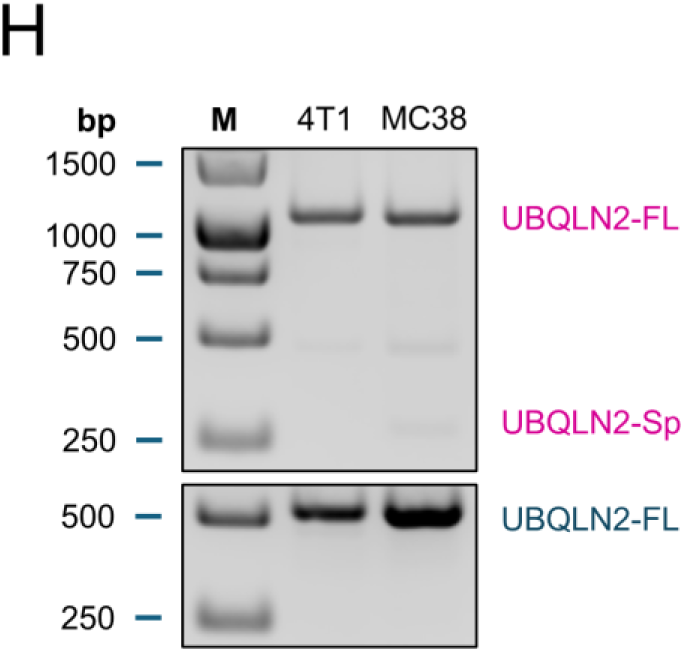
*UBQLN2* contains an unannotated exitron. (A) PacBio-UHRR and ONT-SGNex-HepG2 data as viewed on UCSC genome browser. Selected full-length reads were presented as *horizontal bars* when aligned to *UBQLN2* reference sequence (RefSeq). *Lines* represent regions that are missing in the reads of spliced UBQLN2 mRNAs. Spliced reads share the same SD and were annotated with *red* (SA1) and *black arrows* (SA2). OMIM alleles track indicates the positions of the four codons within the PRR domain (P497, P506, P509, and P525) that are linked to ALS15. UBQLN2 protein domains as translated from the UBQLN2 RefSeq were shown in the alignment. Both spliced UBQLN2 transcripts were also identified from Illumina-based short-read RNA sequencing datasets from TargetALS (ExternalSampleID: CGND-HRA-00013; ExternalSubjectID: NEUEL133AK6) by reads spanning the respective splice junctions and annotated with *red* and *black arrows*. (B) PCR and qPCR primer designs with a shared 5’ primer (*dark blue*) and two alternative 3’ primers to amplify only the full-length (*dark blue*) or both the full-length and spliced UBQLN2 transcripts (*magenta*). (C) Spliced UBQLN2 mRNA (UBQLN2-Sp) was detected in three immortalized human cell lines (U-2 OS, HeLa, and HEK293T) as well as a commercially acquired human brain (HsBrain) RNA sample (Invitrogen AM7962). Note that full-length UBQLN2 (UBQLN2-FL) amplification by the *magenta* splicing primer is inefficient due to competition from the smaller UBQLN2-Sp amplicons (top panel). The bottom panel shows selective amplification of UBQLN2-FL using *dark blue* reverse primer. (D) Percent Spliced Out (PSO) of *HsUBQLN2* exitron in U-2 OS, HeLa, HEK293T, and HsBrain samples as estimated by qPCR splicing assay. Bar height corresponds to mean ± SEM. Each dot represents a biological replicate, N = 4 for U2-OS; N = 5 for HeLa; and N = 3 for HEK293T. (E) DNA sequencing chromatogram from the indicated UBQLN2-Sp RT-PCR amplicons demonstrating the removal of the exitron at the SD-SA2 splice junction marked by the *magenta inverted arrow*. (F) Representative agarose gel image of *UBQLN2* splicing assay from C57BL/6J mouse brain tissues suggests *MmUBQLN2* splicing occurs less frequently. (G) PSO of *MmUBQLN2* exitron estimated from qPCR assay. Four brain regions (right cortex, left cortex, hippocampus, and cerebellum) were harvested and processed from three males (M1, M2, and M3) as well as a female mouse (M4). Bar height corresponds to mean ± SEM. Each dot represents a brain region from an individual mouse as listed above. (H) UBQLN2-Sp transcript was not detected in murine mammary carcinoma 4T1 and colon adenocarcinoma MC38 cells, suggesting that *UBQLN2* exitron splicing is not specific for cancer cells.

We designed an RT-PCR and a qRT-PCR splicing assay using the primer design illustrated in Fig. 1B: a shared forward primer and two different reverse primers to amplify both the full-length (UBQLN2-FL) and spliced UBQLN2 (UBQLN2-Sp) transcripts in human and mouse samples. In all three immortalized human cell lines that we screened, namely U-2 OS, HeLa, and HEK293T, we identified the presence of UBQLN2-Sp transcript from the usage of SA2 (Fig. 1C-E). UBQLN2-SA2 transcript was also detected in a commercially available human total brain RNA sample (Invitrogen AM7962, 30 years old female Caucasian) (Fig. 1C,D; HsBrain), and human spinal cord RNA from both control and ALS patients (not shown). The percent spliced out (PSO) of *HsUBQLN2* exitron in human samples was estimated to be ∼10% by qRT-PCR (Fig. 1D). The identity of the UBQLN2-Sp amplicon from Fig. 1C was validated with Sanger sequencing to be UBQLN2-SA2 (Fig. 1E). UBQLN2-SA1 was not detected in any of the human samples we screened by PCR or qPCR amplification.

Based on protein sequence alignment, the corresponding mm10 coordinates for mouse *UBQLN2* (*Mus musculus UBQLN2/MmUBQLN2*) exitron are chrX:153,498,336-153,499,116. Interestingly, we failed to identify any UBQLN2-Sp transcript variants from mouse long-read sequencing data or mouse brain RNA extracted in-house from different brain regions (cortex, cerebellum, and hippocampus) of C57BL/6J strain by a similar PCR and qPCR strategy (Fig. 1F), suggesting the exitron splicing of *MmUBQLN2* is rare. Consistent with this, the PSO of *MmUBQLN2* was estimated to be less than 0.1% by qRT-PCR splicing assay from four mice (M1 – M4) (Fig. 1G). We also screened two immortalized murine cell lines, namely 4T1 (mammary carcinoma) and MC38 (colon adenocarcinoma), which did not show detectable UBQLN2-Sp transcripts (Fig. 1H). These results suggest that *UBQLN2* splicing occurs at very low levels in the mouse.

### PTBP1 and SRSF1 bind to *HsUBQLN2* pre-mRNA to inhibit splicing

The annotated *HsUBQLN2* 5’UTR (NM_013444.4) contains four polypyrimidine tracts (Py 1-4), that are only partially conserved in *MmUBQLN2* (NM_018798.2) (Fig. 2A), despite an overall homology of greater than 60% ^57,93^. These Py tracts constitute putative binding sites for PTBP1 and PTBP2 – RBPs that are well known to regulate alternative splicing in neurons during differentiation (Fig. 2A). We also identified two (purine-rich tracts; Pu1 and Pu2) representing putative SRSF1 binding motifs in the *HsUBQLN2* 5’UTR (Fig. 2A). Pu1 (CCGG**AGGA**CC) ^94^ is disrupted by two G insertions flanking the GGAGGA motif whereas Pu2 (CGG**AGGA**GGCCC) has two point mutations from AGGA to AGAC in *MmUBQLN2* (Fig. 2A). Py and Pu motifs are also present in the primate *UBQLN2* mRNA sequences (*Pan troglodytes* (XM_001148687.7), *Pongo abelii* (XM_002831718.6), and *Pongo pygmaeus* (XM_054472304.2) (data not shown).

**Fig. 2.**
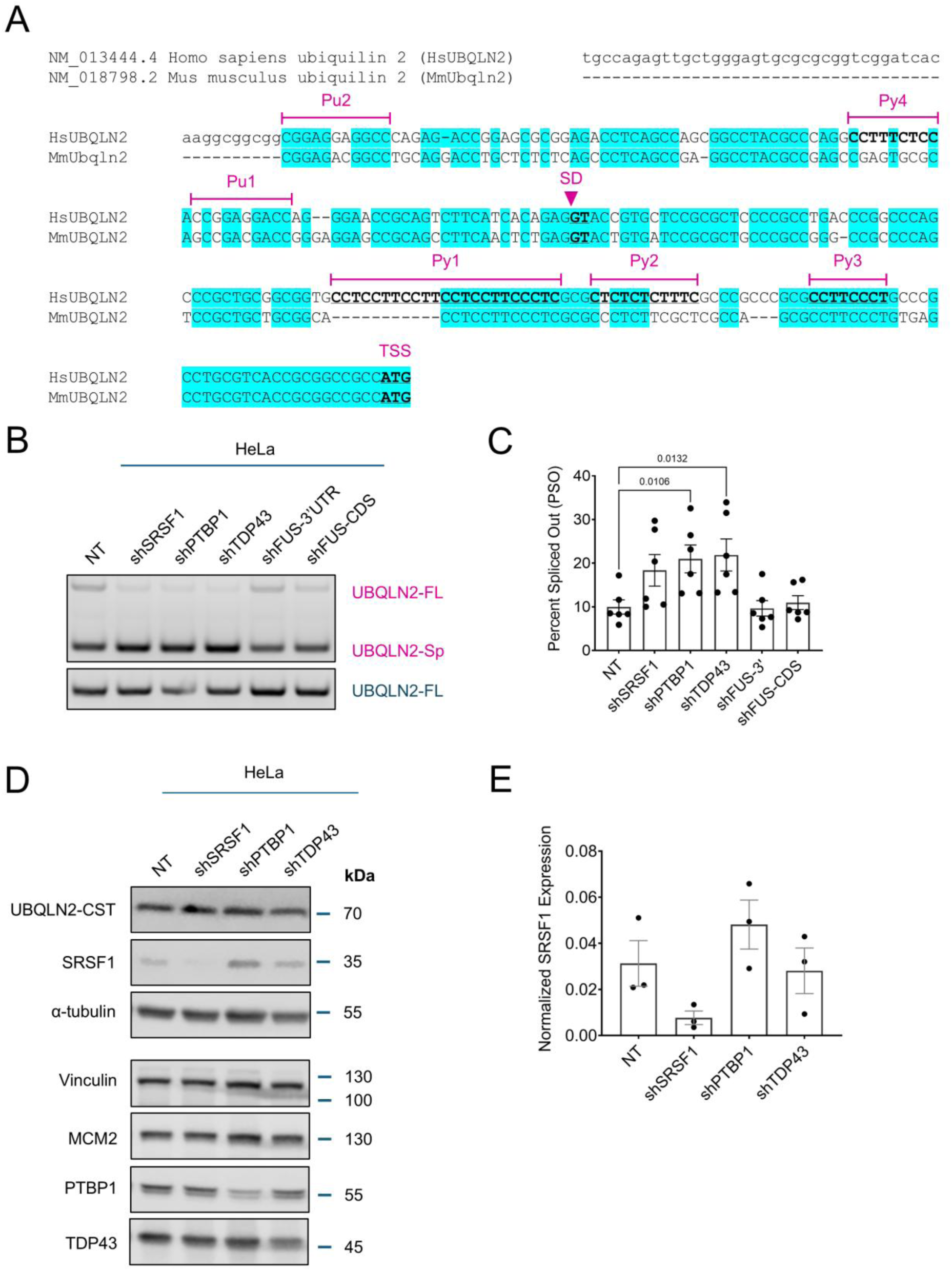
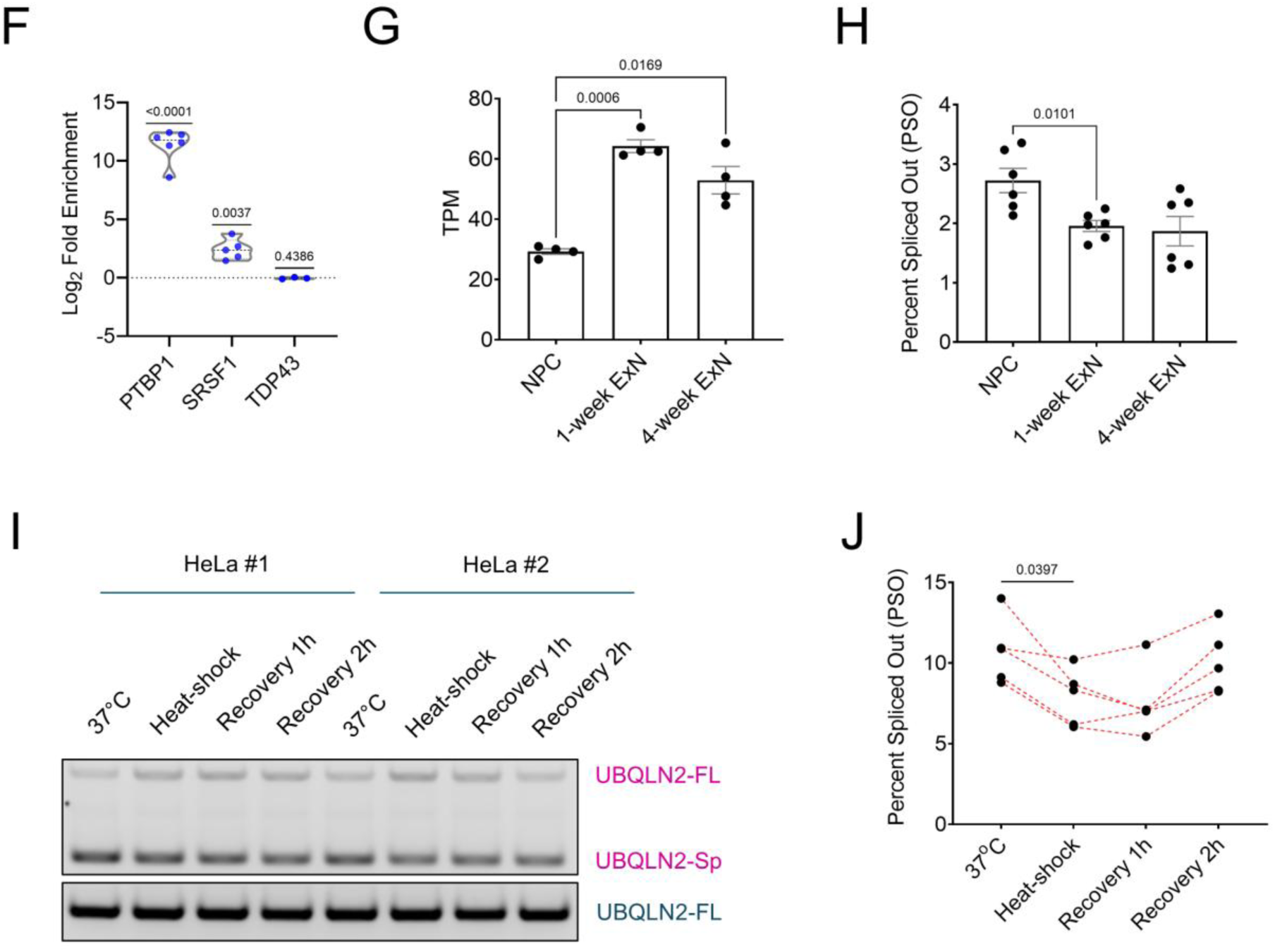
*HsUBQLN2* exitron splicing is inhibited by PTBP1 and SRSF1. (A) Alignment of *UBQLN2* mRNA sequence around the SD (*inverted magenta arrow*) in the 5’ UTR and the translational start site (TSS) of *Homo sapiens* (NM_013444.4) and *Mus musculus* (NM_018798.2) downloaded from NCBI on 15-Sep-2025. Four polypyrimidine tracts (Py1, chrX:56563793-56563815; Py2, chrX:56563819-56563829; Py3: chrX:56563841-56563848; Py4: 56563695-56563703) and two purine-rich SRSF1 binding sites (Pu1, chrX:56563705-56563714; Pu2: chrX:56563639-56563650) on *HsUBQLN2* are highlighted. All elements are unique to *HsUBQLN2* except Py3. (B) HeLa cells were transduced with lentiviral shRNA vectors targeting the indicated RBPs and expression of UBQLN2-FL and UBQLN2-Sp transcripts measured by RT-PCR. (C) Quantification of UBQLN2 exitron PSO from samples analyzed in panel (B) by qRT-PCR assay. Bar height corresponds to mean ± SEM. Each dot represents a biological replicate, N = 6. Significant *p*-values from repeated measures one-way ANOVA with Geisser-Greenhouse correction and Dunnett’s multiple comparisons test were listed. (D) Western blotting of the protein lysates harvested from lentiviral shRNA-transduced HeLa cells from (B) verifying the knockdown of the PTBP1, SRSF1, and TDP43. α-tubulin, vinculin, and MCM2 were included as loading controls for each gel. (E) SRSF1 expression was quantified based on densitometry from (D) and normalized to α-tubulin. Note that shPTBP1 caused an increase in SRSF1 expression. Each dot represents a biological replicate, N = 3. (F) Log_2_ fold enrichment of UBQLN2 pre-mRNA immunoprecipitated from the indicated antibody after normalized to normal IgG control from native RNA immunoprecipitation assays in HeLa cells. Median and interquartile range were shown as dotted lines. Each dot represents an individual biological replicate, N = 6 for PTBP1 RNA-IP, N = 5 for SRSF1 RNA-IP, and N = 3 for TDP43 RNA-IP. The *p*-values from two-tailed one sample t-test were listed. (G) Transcript per million (TPM) of *UBQLN2* from neural progenitor cells (NPC), 1-week differentiating excitatory neurons (1-week ExN), and 4-week differentiating ExN (4-week ExN). Bar height corresponds to mean ± SEM. Each dot represents a biological replicate, N = 2 for a total of two cell lines: GM00498-4 and WC032i-6007-1; n = 4 differentiation batches. Significant *p*-values from repeated measures one-way ANOVA with Geisser-Greenhouse correction and Tukey’s multiple comparisons test were listed. (H) PSO of *HsUBQLN2* from (G) as estimated by qRT-PCR. Bar height corresponds to mean ± SEM. Each dot represents a biological replicate, N = 3 for a total of two cell lines: GM00498-4 and WC032i-6007-1, n = 6 differentiation batches. Significant *p*-values from repeated measures one-way ANOVA with Geisser-Greenhouse correction and Tukey’s multiple comparisons test were listed. (I) HeLa cells were exposed to a 42°C heat shock for 2 h and then immediately harvested or allowed to recover for 1-2 h before cell harvest and RT-PCR analysis of *UBQLN2* splicing. Control cells were kept at 37°C throughout the experiment and harvested at the last timepoint. Representative gel images from two biological replicates were shown. (J) Line graphs showing the changes in PSO as estimated by qRT-PCR over the time course of two hours of heat shock at 42°C followed by two hours of recovery at 37°C. Each line represents a biological replicate, N = 4. Significant *p*-value from repeated measures one-way ANOVA with Geisser-Greenhouse correction and Dunnett’s multiple comparisons test was listed.

Given the presence of consensus PTBP and SRSF1 binding sites, we tested the effects of shRNA-mediated silencing of either SRSF1, PTBP1 or ALS-associated TDP-43 and FUS on *HsUBQLN2* splicing. Knockdown of PTBP1 and TDP-43 significantly increased the PSO of *UBQLN2* exitron in HeLa cells, while knockdown of SRSF1 caused a trend toward increased PSO that was just short of statistical significance (Fig. 2B,C, Sup. Fig. 1A). FUS knockdown did not change the PSO (Fig. 2B,C, Sup. Fig. 1A). The knockdown efficiencies of PTBP1, SRSF1, and TDP-43 shRNAs were confirmed by Western blot (Fig. 2D), where it was also observed that PTBP1 knockdown mildly increased SRSF1 expression (Fig. 2D,E).

In light of these splicing results, we employed native RNA-IP to determine if PTBP1, SRSF1, TDP-43, associated with *UBQLN2* pre-mRNA. As shown in Fig. 2F, both PTBP1 and SRSF1 were significantly enriched on *UBQLN2* pre-mRNA while TDP-43 was not. *STMN2* amplification was included as a positive control for TDP-43 enrichment (Sup. Fig. 1B). These findings suggest that PTBP1 and SRSF1 suppress *UBQLN2* exitron splicing through direct binding, whereas TDP-43 indirectly suppresses *UBQLN2* splicing.

As PTBPs are involved in alternative splicing regulations in response to neuronal differentiation ^35,41–43^, and Ubqln2 expression is upregulated during mouse embryonic stem cell differentiation into neurons ^95^, we examined the total *UBQLN2* gene expression (transcript per million/TPM, Fig. 2G) ^90^ as well as the PSO of *UBQLN2* exitron in human neural progenitor cells (NPC), immature excitatory neurons (ExNs) at 1-week differentiation (1-week ExN), and at 4-week differentiation (4-week ExN) (Fig. 2H). The upregulation in *UBQLN2* gene expression in 1-week neurons compared to NPCs (Fig. 2G) was associated with higher expression of PTBP2 ^96^ and lower PSO (Fig. 2H). To determine whether *UBQLN2* splicing was responsive to proteotoxic stress, we performed

RT-PCR splicing assay in HeLa cells exposed to a transient heat shock. We found that heat shock significantly suppressed UBQLN2 splicing and led to an increase in UBQLN2-FL transcripts (Fig. 2I,J). These findings suggest that *UBQLN2* exitron splicing is regulated and that suppression of exitron splicing may provide a mechanism to upregulate *UBQLN2* gene dosage in response to neuronal differentiation signal and heat-shock.

### Minigenes recapitulate endogenous *UBQLN2* splicing in cell lines

To better understand regulation of *UBQLN2* exitron splicing in human and mouse, we cloned the 5’UTR and the CDS of *HsUBQLN2* and *MmUBQLN2* into a peGFP-N1 vector to generate a *UBQLN2* minigene splicing reporter transcribed from a CMV promoter that produces transcripts and proteins which are tagged with eGFP at the 3’ end and C-terminus, respectively (Fig. 3A). The splicing reporters were transfected into either HeLa or HEK293T cells and RT-PCR splicing analysis was carried out using *UBQLN2* reverse primers (Fig. 1B) in conjunction with a vector-specific forward primer to selectively amplify the minigene-derived transcripts (Fig. 3A). The *HsUBQLN2* minigene showed robust exitron splicing (Fig. 3B,C) with UBQLN2-Sp accounting for 35.8% and 26.8% of total *UBQLN2* mRNA in HeLa and HEK293T cells, respectively (Fig. 3F). By contrast, the mean PSO for *MmUBQLN2* was only 3.9% and 0.8% in these cell lines (Fig. 3D-F). These data establish *UBQLN2* minigenes as reporters of *UBQLN2* splicing activity and are consistent with the finding of dramatically reduced endogenous *MmUBQLN2* splicing in C57BL/6J mouse samples relative to *HsUBQLN2* splicing in human samples (Fig. 1).

**Fig. 3.**
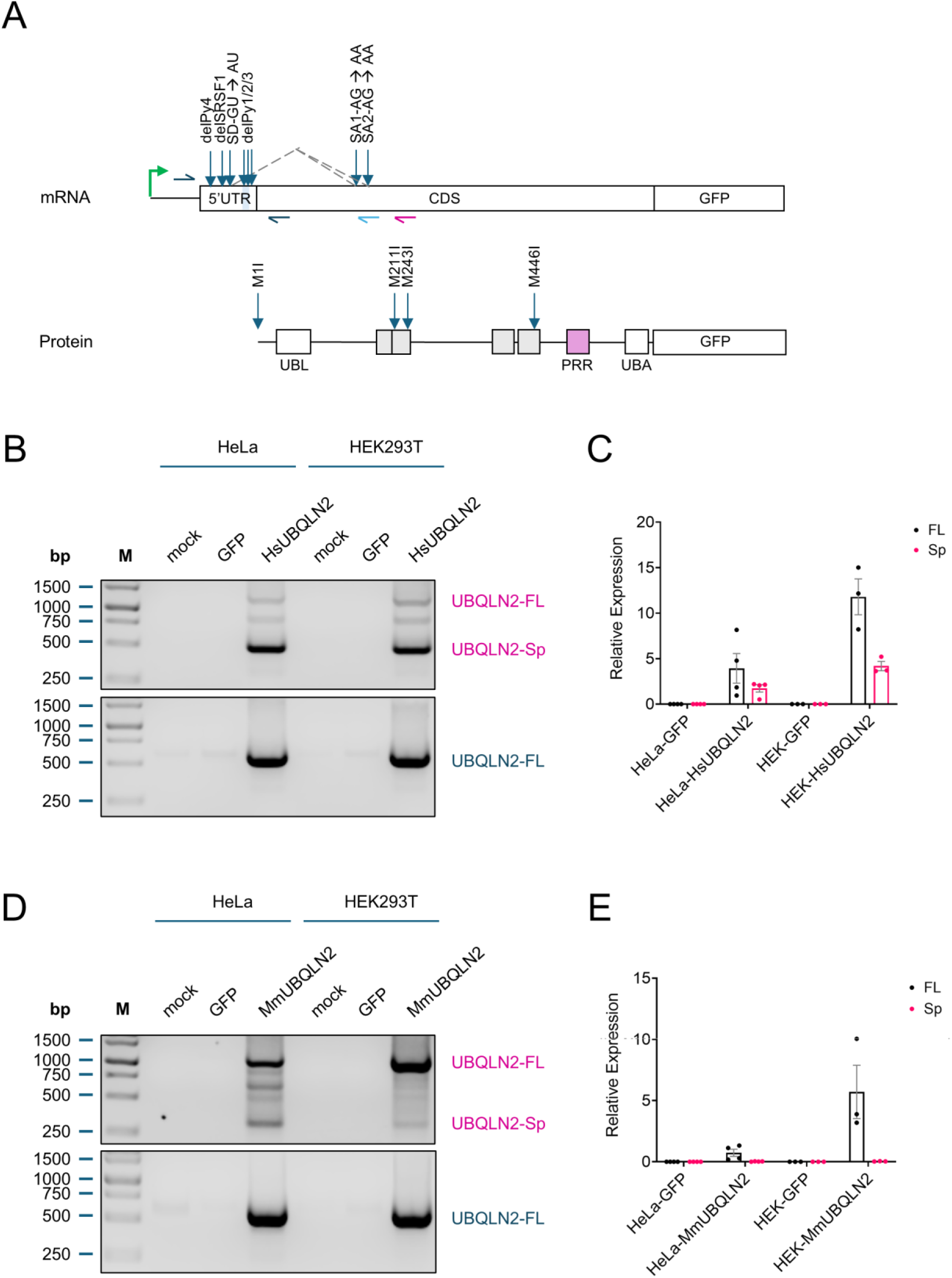
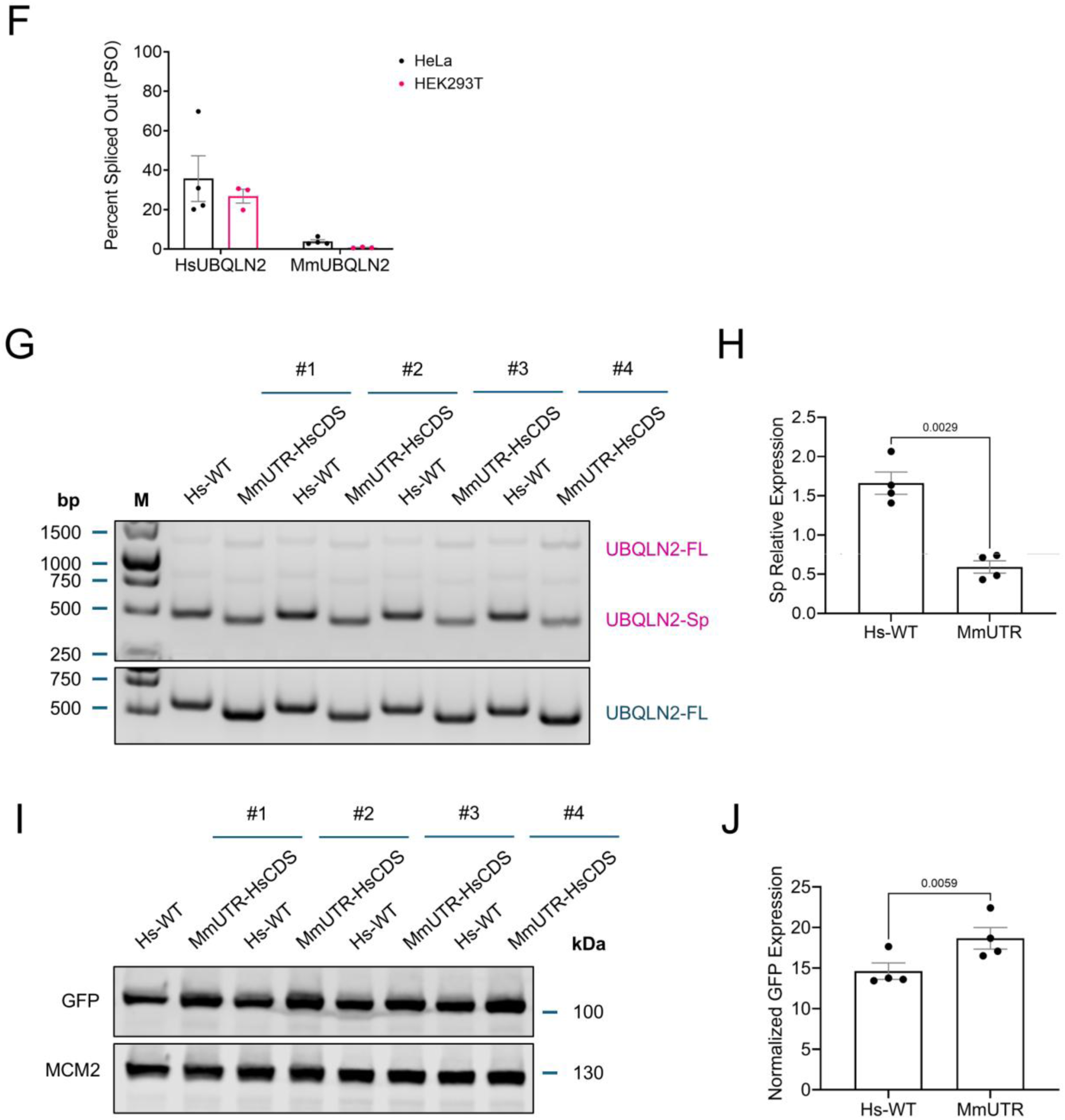
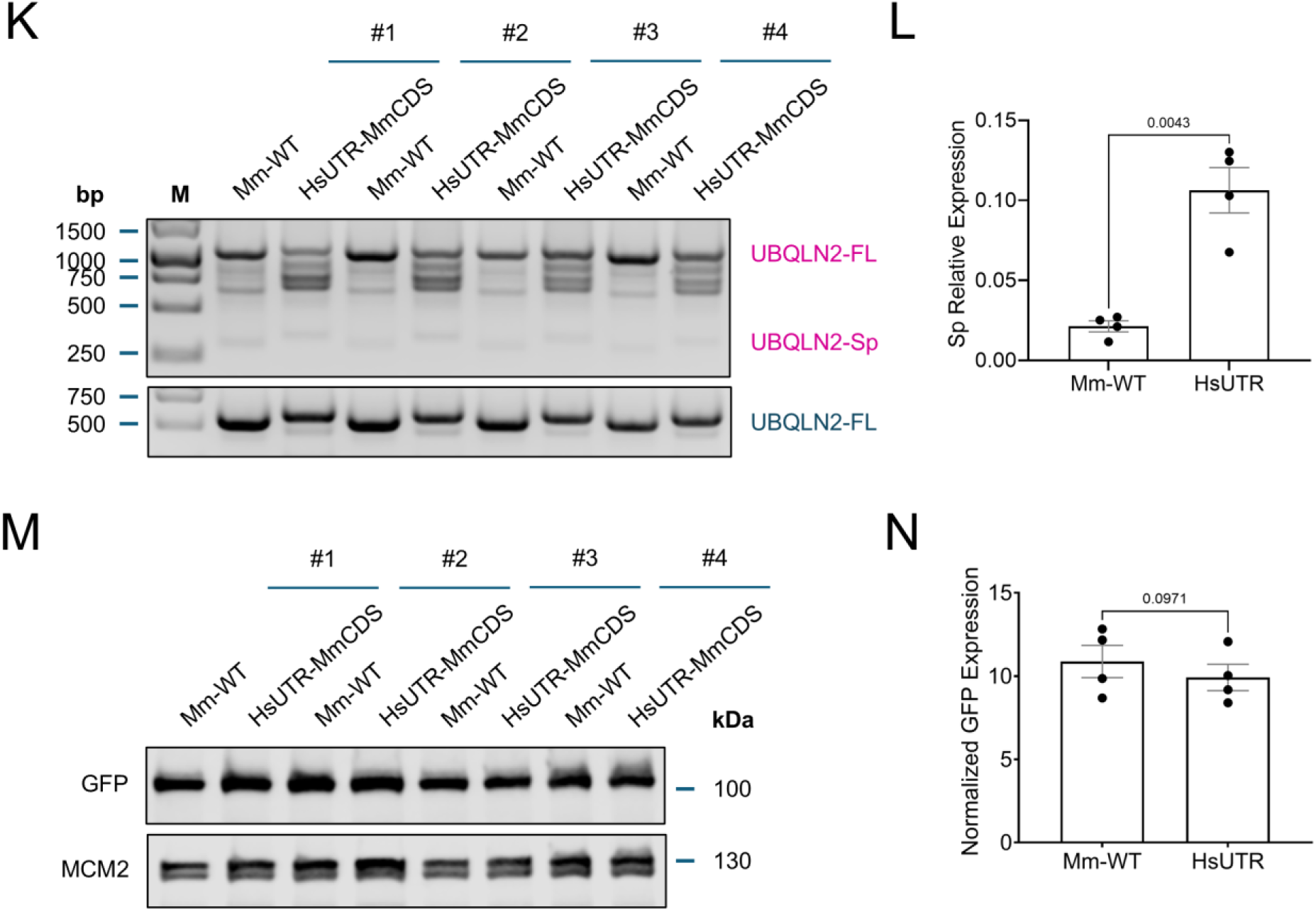
Development of *UBQLN2* minigene splicing reporters. (A, *top* panel) Schematic of the *UBQLN2* splicing reporters. Transcription start site (TSS) from CMV promoter was marked by *green arrow*. Forward RT-PCR primer is located in the multiple cloning site of the vector. Reverse primer between SA1 and SA2 that amplifies mRNAs that are spliced upstream of SA2 is shown in *light blue*. Locations of selected splicing elements and their respective mutants are shown. (A, *bottom* panel) Approximate locations of the TSSs that were mutated in the Hs*UBQLN2* splicing reporter. All primers used for mutagenesis are listed in Sup. Table 2. (B) *HsUBQLN2* splicing reporters were transfected into either HeLa or HEK293T cells. *UBQLN2* splicing assay was performed. (C) The expression of UBQLN2-FL and UBQLN2-Sp transcripts in *HsUBQLN2* transfected cells were quantified by qRT-PCR and normalized to GAPDH. Bar height corresponds to mean ± SEM. Each dot represents a biological replicate, N = 4 for HeLa and N = 3 for HEK293T. (D) *MmUBQLN2* splicing reporters were transfected into either HeLa or HEK293T cells. *UBQLN2* splicing assay was performed. Note that *MmUBQLN2* splicing reporter has negligible UBQLN2-Sp amplicons in both HeLa and HEK293T compared to *HsUBQLN2* as shown in panel (B). (E) The expression of UBQLN2-FL and UBQLN2-Sp transcripts in *MmUBQLN2* transfected cells were quantified by qRT-PCR and normalized to GAPDH. Bar height corresponds to mean ± SEM. Each dot represents a biological replicate, N = 4 for HeLa and N = 3 for HEK293T. represents a biological replicate, N = 4 for HeLa and N = 3 for HEK293T. (G) The 5’UTR of *HsUBQLN2* reporter was replaced with the annotated 5’UTR from *MmUBQLN2* (MmUTR-HsCDS). Splicing assay of both Hs-WT and MmUTR-HsCDS reporter-transfected cells was shown. (H) MmUTR replacement reduced the relative expression of UBQLN2-Sp transcripts, as quantified by qRT-PCR. Bar height corresponds to mean ± SEM. Each dot represents a biological replicate, N = 4. *p*-value from two-tailed paired t-test was listed. (I) Western blot of UBQLN2-GFP and MCM2 (loading control) from the protein lysates of the transfected cells used in Panel (G). (J) UBQLN2-GFP signal intensity was quantified by densitometry and normalized to MCM2. Bar height corresponds to mean ± SEM. Each dot represents a biological replicate, N = 4. *p*-value from two-tailed paired t-test was listed. (K) The 5’UTR of *MmUBQLN2* reporter was replaced with the annotated 5’UTR from *HsUBQLN2* (HsUTR-MmCDS). Splicing assay of both Mm-WT and HsUTR-MmCDS reporter-transfected cells was shown. (L) HsUTR replacement increased the relative expression of UBQLN2-Sp transcripts as quantified by qRT-PCR. Bar height corresponds to mean ± SEM. Each dot represents a biological replicate, N = 4. *p*-value from two-tailed paired t-test was listed. (M) Western blot of UBQLN2-GFP and MCM2 (loading control) from the protein lysates of the transfected cells listed in Panel (K). (N) UBQLN2-GFP signal intensity was quantified by densitometry and normalized to MCM2. Bar height corresponds to mean ± SEM. Each dot represents a biological replicate, N = 4. *p*-value from two-tailed paired t-test was listed.

The relative absence of exitron splicing of *MmUBQLN2* in human cells suggests the presence of unique *cis*-acting splicing elements between *HsUBQLN2* and *MmUBQLN2* genes. Consistent with this idea, substitution of the HsUBQLN2 5’UTR (HsUTR) with the MmUBQLN2 5’UTR (MmUTR) (see Materials and Methods) decreased splicing of the *HsUBQLN2* minigene reporter approximately three-fold (Fig. 3G,H). The decrease in splicing of the MmUTR-HsUBQLN2 chimera correlated with a small but statistically significant increase in HsUBQLN2-M1 protein expression (Fig. 3I,J). We also performed the converse experiment where the effect of the HsUTR on the splicing of the *MmUBQLN2* reporter was measured. The HsUTR-MmUBQLN2 chimeric construct exhibited a complex splicing pattern characterized by the presence of multiple spliced products that were shorter than the canonical MmUBQLN2-FL transcript (Fig. 3K). We postulate that these transcripts are produced through usage of the stronger SD site located within the HsUTR which has the tendency to be spliced with multiple weaker downstream SA sites in *MmUBQLN2*. In support of the idea that the HsUTR contains elements that enhance splicing, the HsUTR-MmUBQLN2 chimera showed ∼5-fold increased relative expression of spliced MmUBQLN2 transcript (Fig. 3K,L). However, the fraction of HsUTR-MmUBQLN2-Sp (Fig. 3L) was still 5-10-fold less than what was observed for HsUBQLN2-Sp (Fig. 3H). MmUBQLN2-M1 protein expression showed a slight reduction correlating with the increase in HsUTR-MmUBQLN2 splicing (Fig. 3M,N). These findings suggest the presence of *cis*-acting splicing elements unique to the *HsUBQLN2* 5’UTR to regulate its exitron splicing and expression. However, *cis*-acting elements outside the 5’UTR also contribute to splicing differences between *MmUBQLN2* and *HsUBQLN2*.

### Identification of *cis*-acting elements that modulate *UBQLN2* splicing

We next sought to identify *cis*-acting elements that specify *HsUBQLN2* exitron removal by introducing silent mutations into the SD (GU to AU) and the two alternative SAs (AG to AA) (Fig. 3A). We also deleted four Py tracts and two purine-rich, putative SRFS1 binding motifs (Pu1-2), or introduced point mutations within these motifs, to investigate the roles PTBP1 and SRSF1 in *HsUBQLN2* exitron splicing (Fig. 3A,4A). Mutation of the putative SD (mean PSO: 0.2%) completely inhibited *HsUBQLN2* exitron splicing (WT mean PSO: 31.6%) while mutation of SA2 inhibited most splicing events with a residual mean PSO of ∼2% (Fig. 4B,C, Sup. Fig. 1C). To test whether residual splicing of the SA2 mutant was due to usage of upstream SA1, we measured splicing of a *HsUBQLN2* SA1/SA2 double mutant. Splicing was still detected in this mutant with a mean PSO of 2.4%, suggesting that an alternative upstream SA can be used in the absence of SA1 and SA2 (Fig. 4B,C). To detect transcripts using an upstream SA other than SA2, we further designed an additional splicing primer between SA1 and SA2 (Fig. 3A, light blue reverse primer) which produced a significantly higher mean PSO in the SA mutants (∼1% for both mutants versus 0.35% in WT) (Fig. 4D), confirming the residual exitron splicing was a result from the usage of an unidentified upstream SA. We also investigated the effects of mutating the canonical translation start codon (M1) and several candidate start codons downstream from SA2 that may be used to translate UBQLN2-Sp transcripts (Fig.3A). In general, mutating the AUG codons did not change the PSO of *HsUBQLN2* exitron splicing, except a M1/243I double mutant which showed a significant reduction in mean PSO (15.4%) compared to WT (Fig. 4C). Because *MmUBQLN2* splicing reporter produced several rare, spliced transcripts in human cells (Fig. 3D), we decided to also mutate the SD of *MmUBQLN2* to examine its effect on exitron splicing. *MmUBQLN2*-SD mutant was found to eliminate the rare MmUBQLN2-Sp amplicons and reduced the PSO from 5.76% in WT to 0.9% in the SD mutants (Fig. 4E,F).

**Fig. 4.**
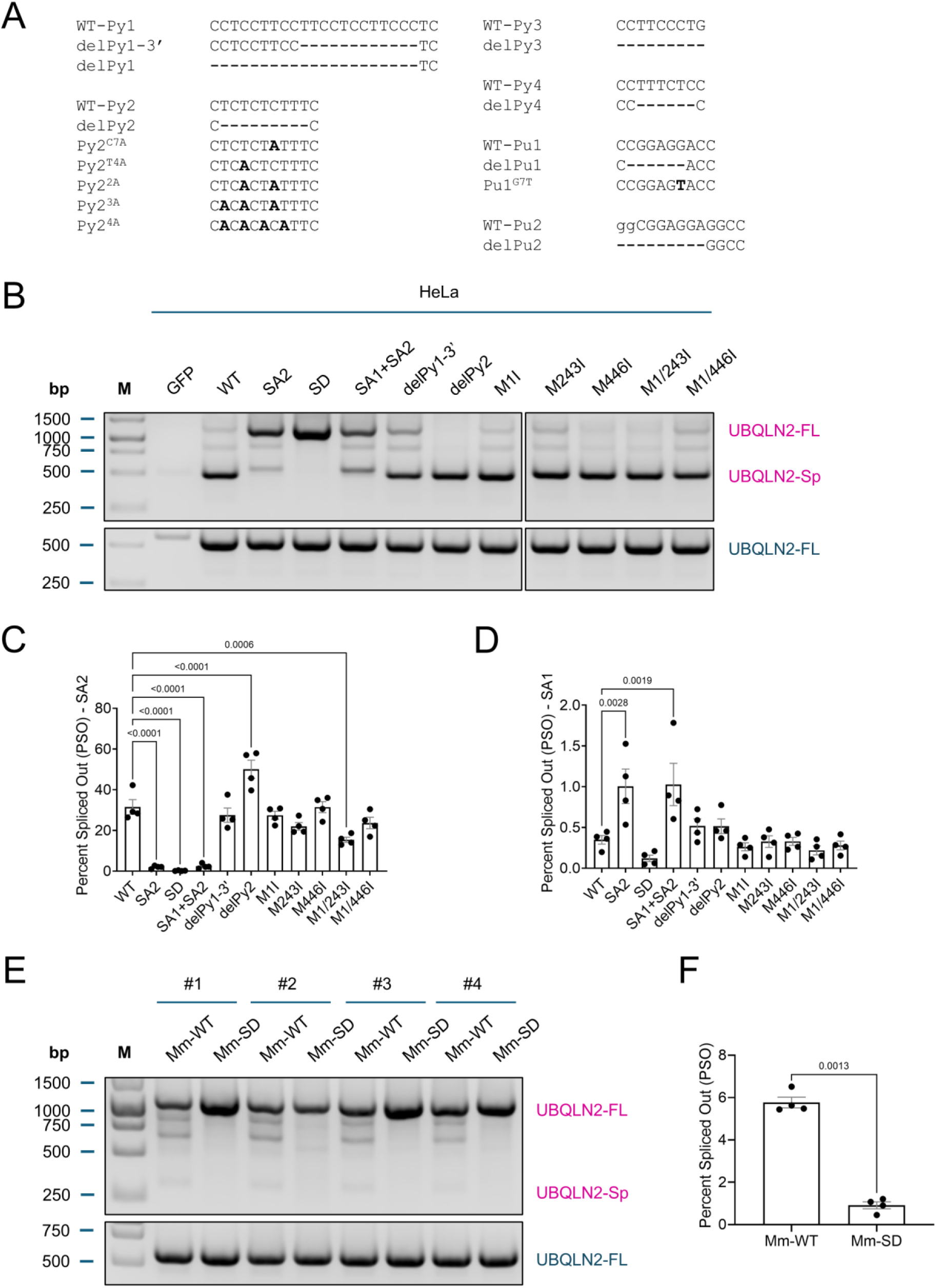
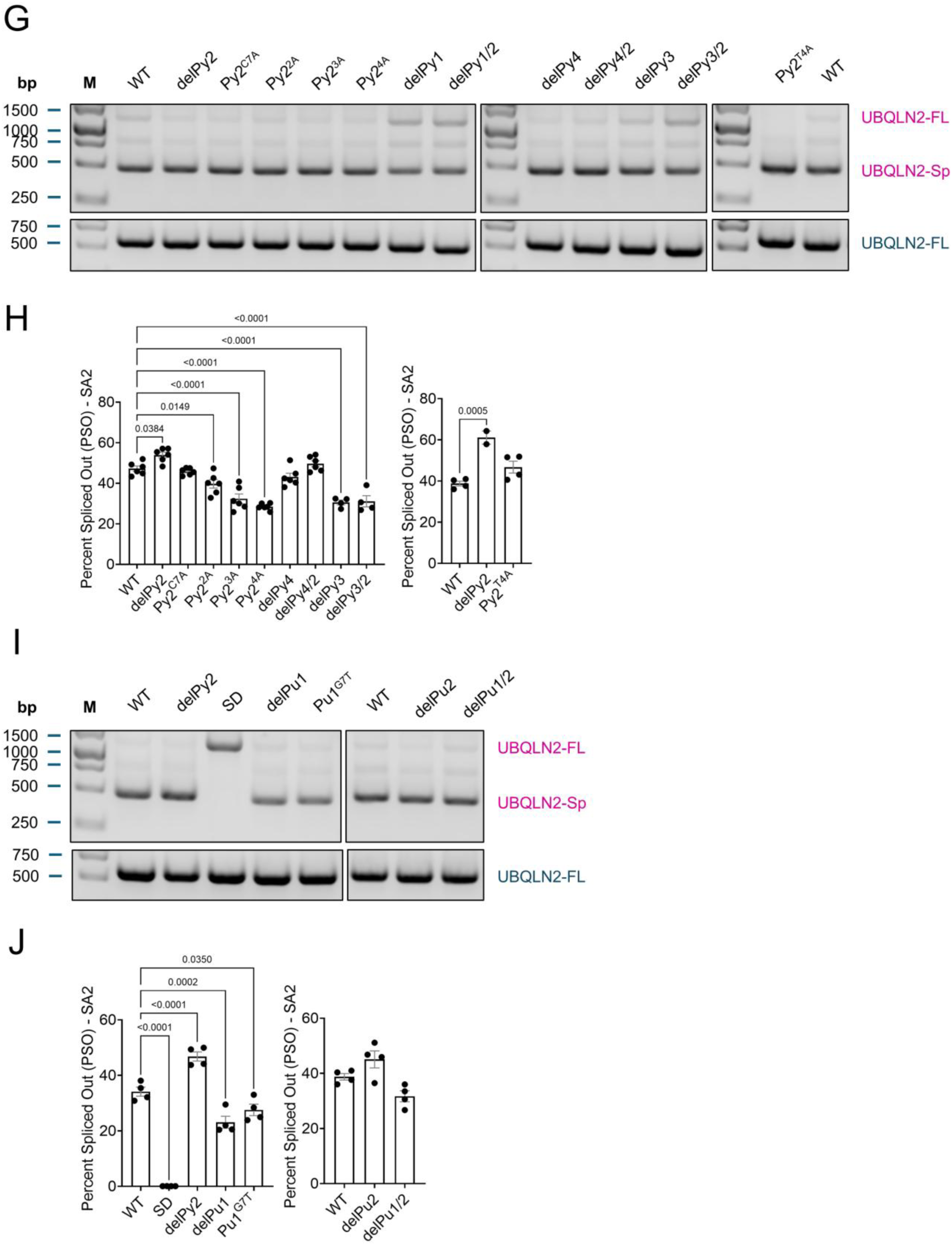
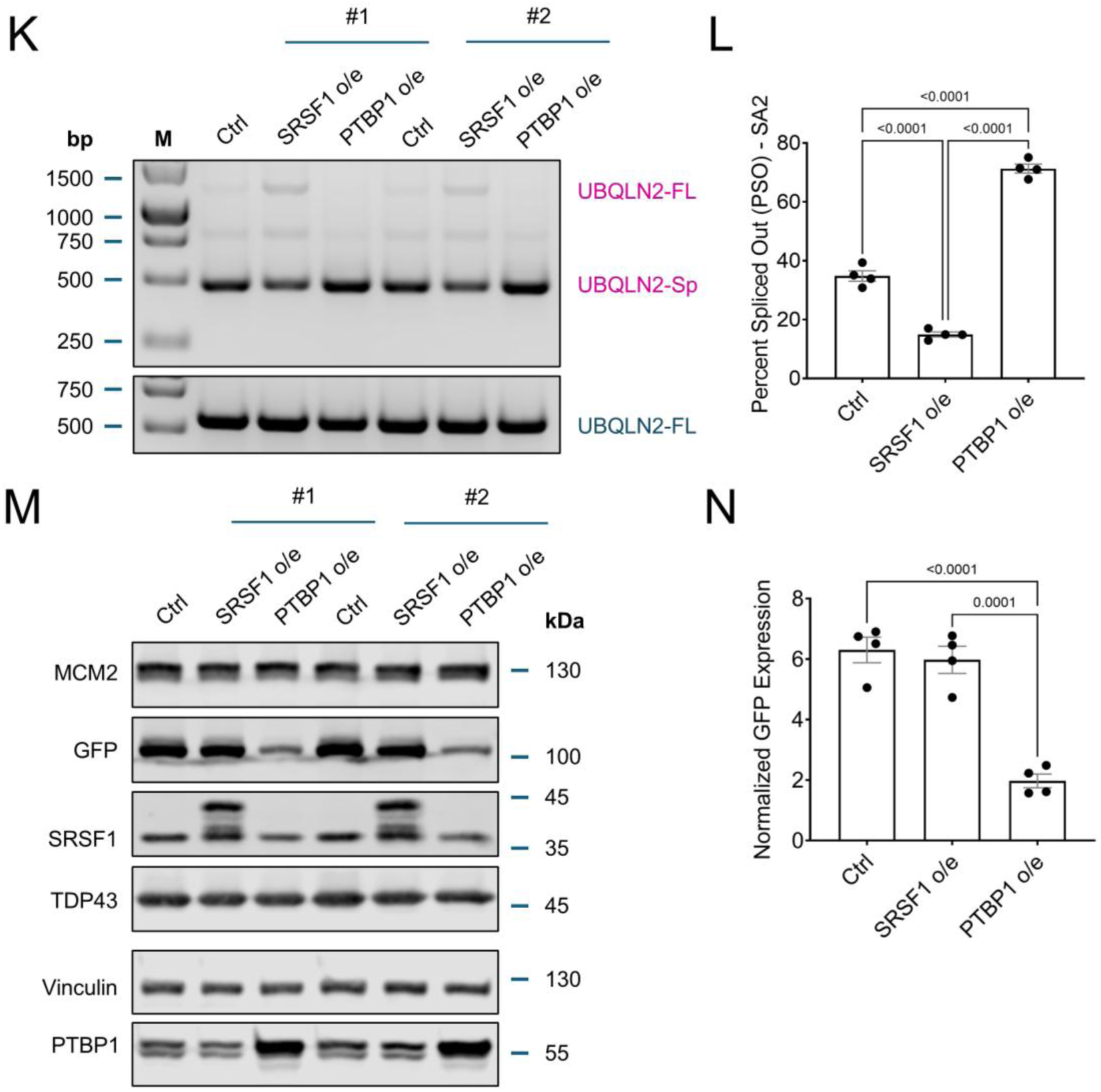
*HsUBQLN2* is spliced from an essential SD and at least two alternative SAs. (A) Py1-4 and Pu1-2 tracts were either deleted or mutated by nucleotide substitutions as shown in *bold*. (B) RT-PCR splicing assay of HeLa cells transfected with wild-type or mutant *UBQLN2* splicing reporters using primers shown in Fig. 3A. Note the absence of the UBQLN2-Sp amplicons in the SD mutant and an upward shift of the UBQLN2-Sp amplicons in the SA2 and SA1+SA2 double mutants, suggesting the usage of an upstream unidentified SA. (C, D) Quantification of UBQLN2 PSO-SA2 and PSO-SA1 in HeLa cells expressing the indicated *UBQLN2* minigenes by qRT-PCR. Bar height corresponds to mean ± SEM. Each dot represents a biological replicate, N = 4, significant *p*-values from one-way ANOVA and Dunnett’s multiple comparisons test were listed. (E) RT-PCR splicing assay of HeLa cells transfected with wild-type or mutant *MmUBQLN2*-WT or *MmUBQLN2*-SD minigenes. Note the disappearance of the UBQLN2-Sp amplicons and the increase in intensity of the UBQLN2-FL amplicons, N = 4. (F) Quantification of the PSO from (E) by qRT-PCR. Bar height corresponds to mean ± SEM. Each dot represents a biological replicate, N = 4, *p*-value from two-tailed one sample *t*-test was listed. (G) RT-PCR splicing assay of HeLa cells transfected with splicing reporters harboring mutations in Py motifs shown in Fig. 4A. (H) Quantification of PSO-SA2 of panel (G) by qRT-PCR. Bar height corresponds to mean ± SEM. Each dot represents a biological replicate, 6 ≥ N ≥ 4, significant *p*-values from one-way ANOVA and Dunnett’s multiple comparisons test were listed. Note that the delPy1 mutation disrupted primer binding and levels of UBQLN2-FL products harboring this mutation could not be determined by qPCR. (I) RT-PCR splicing assay of HeLa cells transfected with splicing reporters harboring mutation in Pu tracts as shown in Fig. 4A. (J) Quantification of PSO-SA2 of panel (I) by qRT-PCR. Bar height corresponds to mean ± SEM. Each dot represents a biological replicate, N = 4, significant *p*-values from one-way ANOVA and Dunnett’s multiple comparisons test were listed. (K) *UBQLN2* splicing assay from HeLa cells co-transfected with *HsUBQLN2* splicing reporter and SRSF1 (SRSF1 o/e) or PTBP1 plasmids (PTBP1 o/e). (L) PSO of *HsUBQLN2* from (K) as estimated by qRT-PCR. Bar height corresponds to mean ± SEM. Each dot represents a biological replicate, N = 4. Significant *p*-values from one-way ANOVA and Tukey’s multiple comparisons test are shown. (M) Western blot showing expression of PTBP1 and SRSF1 in transfected HeLa cells used in panel (K). Note the reduction of full-length UBQLN2-GFP and SRSF1 expression in PTBP1 o/e cell lysates. TDP-43 and MCM2 were included as loading controls. (N) UBQLN2-GFP signal intensity was quantified by densitometry and normalized to MCM2. Bar height corresponds to mean ± SEM. Each dot represents a biological replicate, N = 4. Significant *p*-values from one-way ANOVA and Tukey’s multiple comparisons test were listed.

Given that PTBP1 and SRSF1 occupied *HsUBQLN2* mRNA (Fig. 2F) and their knockdown suppressed *HsUBQLN2* splicing (Fig. 2B,C), we tested the effects of deleting the putative PTBP1 (Py1-4) and SRSF1 (Pu1-2) binding motifs in the 5’UTR of the *HsUBQLN2* minigene (Fig. 2A,3A,4A). *HsUBQLN2* Py1 contains a tandem CCTCCTTCC motif that is present as a single copy in MmUBQLN2 (Fig. 2A). Deletion of the 3’ CCTCCTTCC repeat in Py1 (delPy1-3’) (Fig. 4A) did not significantly affect *HsUBQLN2* splicing, however, a deletion of the entire Py1 motif (delPy1) and a double deletion of Py1 and Py2 (delPy1/2) both resulted in an increase in the UBQLN2-FL transcripts accompanied by a reduction in UBQLN2-Sp transcripts (Fig. 4A,G, Sup. Fig. 1D), suggesting that the presence of Py1 promotes *UBQLN2* splicing.

On the other hand, deletion of Py2 phenocopied the enhancing effect of PTBP1 knockdown on endogenous *HsUBQLN2* splicing (Fig. 2B,C), suggesting that PTBP1 binding through Py2 inhibits *HsUBQLN2* splicing (Fig. 4B,C,G,H). By contrast, point mutations in Py2 inhibited UBQLN2 splicing roughly proportional to the number of mutations introduced (Fig. 4A,G,H). Deletion of Py3 (delPy3), which is fully conserved between mouse and human, as well as a double deletion of Py2 and Py3 (delPy3/2) significantly reduced the mean PSO (Fig. 4A,G,H). Deletion of Py4 (delPy4) did not alter the PSO; whereas double deletion of Py4 and Py2 (delPy4/2) slightly increased the PSO (Fig. 4A,G,H). These combined findings suggest that Py2 and Py1/Py3 act antagonistically in *UBQLN2* splicing regulation, with Py2 inhibiting splicing and Py1/Py3 promoting splicing. The net effect of PTBP1 on *UBQLN2* splicing may therefore depend on which sites are occupied and how occupied sites interact with each other.

Deletion (delPu1) and point substitution (Pu1^G7T^) within the putative SRSF1 binding purine track (delPu1) inhibited *HsUBQLN2* exitron splicing, each with lower mean PSO (23.1% and 27.6% respectively) compared to WT (34.1%) (Fig. 4A,I,J). In contrast, deletion of Pu2 track (delPu2) slightly increased the PSO (45.1%) compared to the WT (38.8%) and the trend was reversed with a double deletion of Pu1 and Pu2 (delPu1/2) (Fig. 4A,I,J). These results suggest that the enhancer Pu1 is epistatic to the repressor Pu2. Pu2 might serve as the intronic splicing silencer which is bound by SRSF1 to repress exitron splicing.

We also investigated the effects of overexpressing PTBP1 and SRSF1 on *HsUBQLN2* splicing. SRSF1 overexpression inhibited *HsUBQLN2* exitron splice out (Fig. 4K,L, Sup. Fig. 1E), further establishing SRSF1 as an inhibitor of *HsUBQLN2* exitron splicing in addition to its knockdown (Fig. 2B,C). Surprisingly, overexpression of PTBP1 also increased *UBQLN2* exitron splice out (Fig. 4K,L, Sup. Fig. 1E). Increased splicing correlated with a reduction in HsUBQLN2-M1 protein (Fig. 4M,N). We also observed a slight decrease in SRSF1 expression associated with PTBP1 overexpression (Fig. 4M, Sup. Fig. 1F,G). The fact that both knockdown and overexpression of PTBP1 increased *HsUBQLN2* splicing suggests that *HsUBQLN2* splicing is highly sensitive to PTBP1 gene dosage and/or that PTBP1 influences networks of other RBPs, such as SRSF1, which are direct/indirect regulators of *HsUBQLN2* exitron splicing.

### Alternatively spliced UBQLN2 transcripts produce truncated protein isoforms

Multiple AUG codons downstream of SA2 are in-frame with the HsUBQLN2-FL CDS, raising the possibility that UBQLN2-Sp transcripts could produce N-terminally truncated UBQLN2 proteins (Fig. 5A). Consistent with this, Western blotting of HeLa extracts expressing the *HsUBQLN2* minigene with α-GFP antibodies detected a ∼70 kDa protein (p70) and several lower molecular weight species in addition to ∼100 kDa, full-length UBQLN2-GFP. Fig. 5B, WT). Abundance of the p70 species was reduced by mutation of SA2 or SD (Fig. 3A), suggesting it is produced from the UBQLN2-Sp transcript (Fig. 5B,C). Furthermore, mutation of the canonical UBQLN2 start codon from AUG to AUA (Met to Ile, M1I), increased the expression of p70, suggesting a downstream AUG codon can be used for translation initiation (Fig. 5B,D). We noted that the AUG triplet at codon 243 (M243I) occurred in a more favorable Kozak sequence context compared to the AUG triplet at codon 211 – the first Met downstream of SA2 (Fig. 5A). Indeed, an M243I mutation abolished expression of p70 either with or without the M1I mutation (Fig. 5B,D). Mutation of codon 446 (M446I) had no effect on p70 abundance whereas the M1/446I double mutant showed significantly higher expression of p70, suggesting competitive binding of ribosomes to these suboptimal RBSs (Fig. 5B,D). Finally, we also noticed higher production of a 35 kDa species (p35) in the M1I mutants that could be consistent with translation initiation from Met-446; however, an M446I mutation did not eliminate p35 production (Fig. 5B). These findings suggest that, in addition to UBQLN2-M1, *HsUBQLN2* minigene produces at least one truncated, splicing-dependent, isoform that is initiated at Met-243 (UBQLN2-M243).

**Fig. 5.**
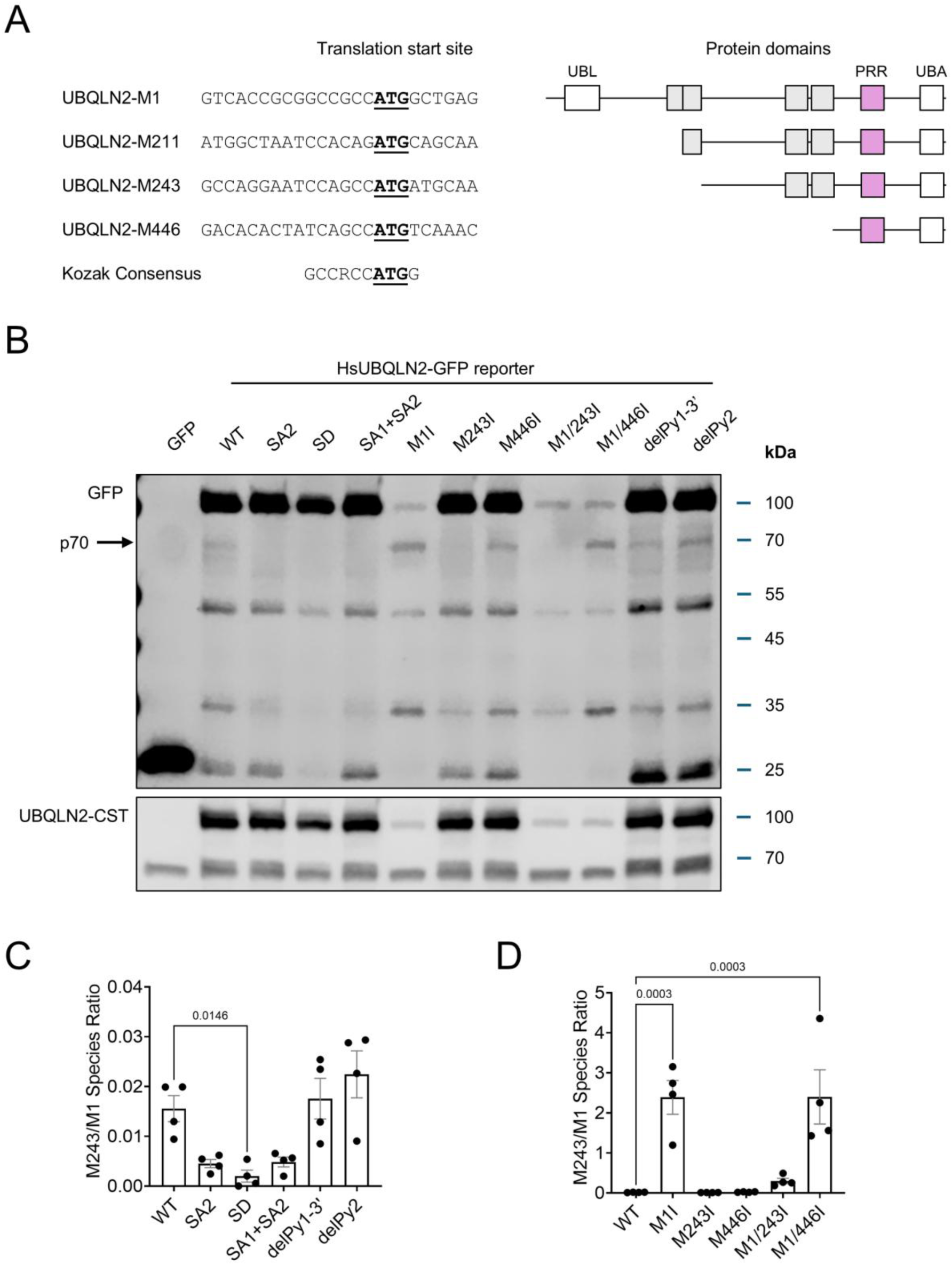
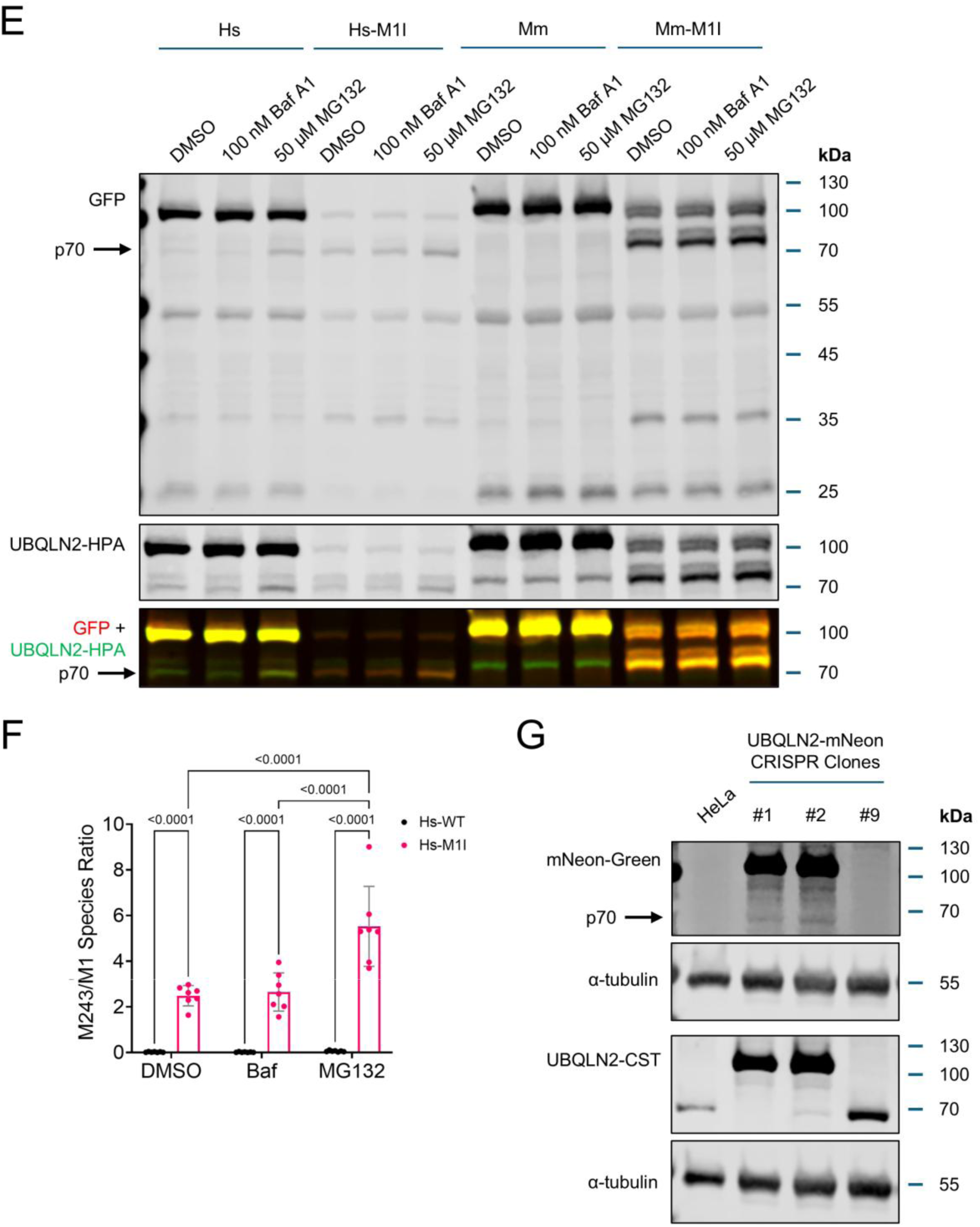
Spliced *HsUBQLN2* is translated from Met-243. (A, *left* panel) Alignment of UBQLN2 start codons downstream of SA2 (*underlined* and in *bold*) with Kozak consensus. (A, *right* panel) Schematic depiction of translational products produced from the indicated AUG codons. (B) Western blotting of the whole cell lysates extracted from HeLa cells transfected with the corresponding mutated *HsUBQLN2* minigene reporters as shown in Fig. 4B with GFP antibody. p70 expression (*black* arrow) is upregulated in M1I mutants and absent in splice site mutants as well as the M243I mutants, indicating it corresponds to UBQLN2-M243. α-UBQLN2-CST (Cell Signaling Technologies 85509S) that detects endogenous UBQLN2 (lower band of the two 70-kDa species) and overexpressed full-length UBQLN2-GFP, but not the N-terminally truncated UBQLN2 species. (C, D) Quantification of the M243/M1 species ratio from (B). Bar height corresponds to mean ± SEM. Each dot represents a biological replicate, N = 4, significant *p*-values from one-way ANOVA and Dunnett’s multiple comparisons test were listed. (E) UBQLN2-M243 accumulates upon proteasome inhibition. HeLa cells were transfected with wild-type *HsUBQLN2* or *MmUBQLN2* reporters or their corresponding M1I mutants. Cells were treated with 100 nM bafA1 or 50 µM MG132 for 4 h and cell lysates were analyzed by Western blotting with the indicated antibodies. p70 species was selectively induced by MG132 treatment. α-UBQLN2-HPA (Sigma HPA006431) that recognizes the C-terminal epitope L443-N590 showed overlapping signal with α-GFP. (F) Quantification of the M243/M1 species ratio from (E). Bar height corresponds to mean ± SEM. Each dot represents a biological replicate, N = 7, significant *p*-values from two-way ANOVA and Tukey’s multiple comparisons test with a single pooled variances were listed. (G) Western blot analysis of unedited HeLa cells and three selected UBQLN2-mNeon CRISPR HeLa clones with α-mNeon-Green antibody. Clones #1 and #2 are positive for mNeon knock-in on at least one allele (Sup. Fig. 3) whereas clones #9 is negative. Vinculin was included as loading control.

The UBQLN2-M243 isoform was expressed at ∼2% of the level of the UBQLN2-M1 isoform in HeLa cells expressing the WT *HsUBQLN2* splicing reporter (Fig. 5C). This relatively low abundance likely reflects the imperfect RBS spanning codon 243 (Fig. 5A). However, reduced stability of UBQLN2-M243 could also contribute to its reduced steady state levels relative to UBQLN2-M1. To explore this, we tested the effects of bafilomycin A1 (BafA, autophagy inhibitor) or MG132 (proteasome inhibitor) on steady state levels of HsUBQLN2-M1 and HsUBQLN2-M243 in HeLa cells expressing either the *HsUBQLN2* WT or M1I minigenes (Fig. 5E,F). Treatment with MG132 stabilized UBQLN2-M243 and doubled the M243/M1 species ratio in cells transfected with either HsUBQLN2-WT or HsUBQLN2-M1I minigenes (Fig. 5E,F). Consistent with the notion that *MmUBQLN2* is weakly spliced, MG132 treatment did not induce a p70-related protein in HeLa cells expressing the MmUBQLN2-WT minigene (Fig. 5E); however, we did detect a ∼75 kDa species in the MmUBQLN2-M1I mutant that likely reflects downstream translation initiation from M254 (equivalent to M243 in *HsUBQLN2*) (Fig. 5E). However, efforts trying to identify the lower molecular weight translation products from MmUBQLN2-M1I mutant failed, as none of the selected Met double mutants (M88, M192, M205, M254, M368, M496, and M606) eliminated the smaller species produced in the MmUBQLN2-M1I mutant (Sup. Fig. 2). Finally, to investigate endogenous UBQLN2 isoform production, we used CRISPR-Cas9-mediated homology-directed repair (HDR) to knock-in an in-frame, C-terminal mNeonGreen fluorescent tag into the *UBQLN2* gene locus in HeLa cells (Sup. Fig. 3). The ∼70 kDa species produced from UBQLN2-M243 was detected by mNeon antibody from two independent CRISPR clones (Fig. 5G, mNeon clone #1 and #2). From the combined findings, we conclude that N-terminally truncated HsUBQLN2-M243 isoform can be expressed from both the minigene reporter as well as from the endogenous locus.

### UBQLN2-M243 exhibits altered inclusion dynamics

To investigate potential functional differences amongst UBQLN2 isoforms, we generated N-terminal GFP-tagged HsUBQLN2 expression vectors corresponding to the M1, M211, M243, and M446 isoforms (Fig. 6A). UBQLN2-M211 and UBQLN2-M446 were omitted from downstream studies either owing to their poor Kozak consensus or undetectable expression levels (Fig. 5A,B). Consistent with previous work from our group and others, overexpressed GFP-UBQLN2-M1 formed large, spherical cytosolic inclusions that may reflect liquid-liquid phase separation (LLPS) and/or endolysosomal localization (Fig. 6B) ^97–100^. By contrast, GFP-UBQLN2-M243 exhibited irregularly shaped structures that were frequently clustered around the nucleus (Fig. 6B). GFP-UBQLN2-M243 inclusions were significantly less round than those formed by GFP-UBQLN2-M1 (Fig. 4C, Sup. Fig. 4A-C), suggesting altered phase separation characteristics. Those spherical inclusions that formed in GFP-UBQLN2-M243-expressing cells tended to be significantly smaller than aggregates formed by GFP-UBQLN2-M1 (Fig. 6B,D). Irregular aggregates formed by GFP-UBQLN2-M243 could be a direct effect of N-terminal truncation on UBQLN2 protein structure or may reflect changes in the UBQLN2 protein-protein interaction landscape.

**Fig. 6.**
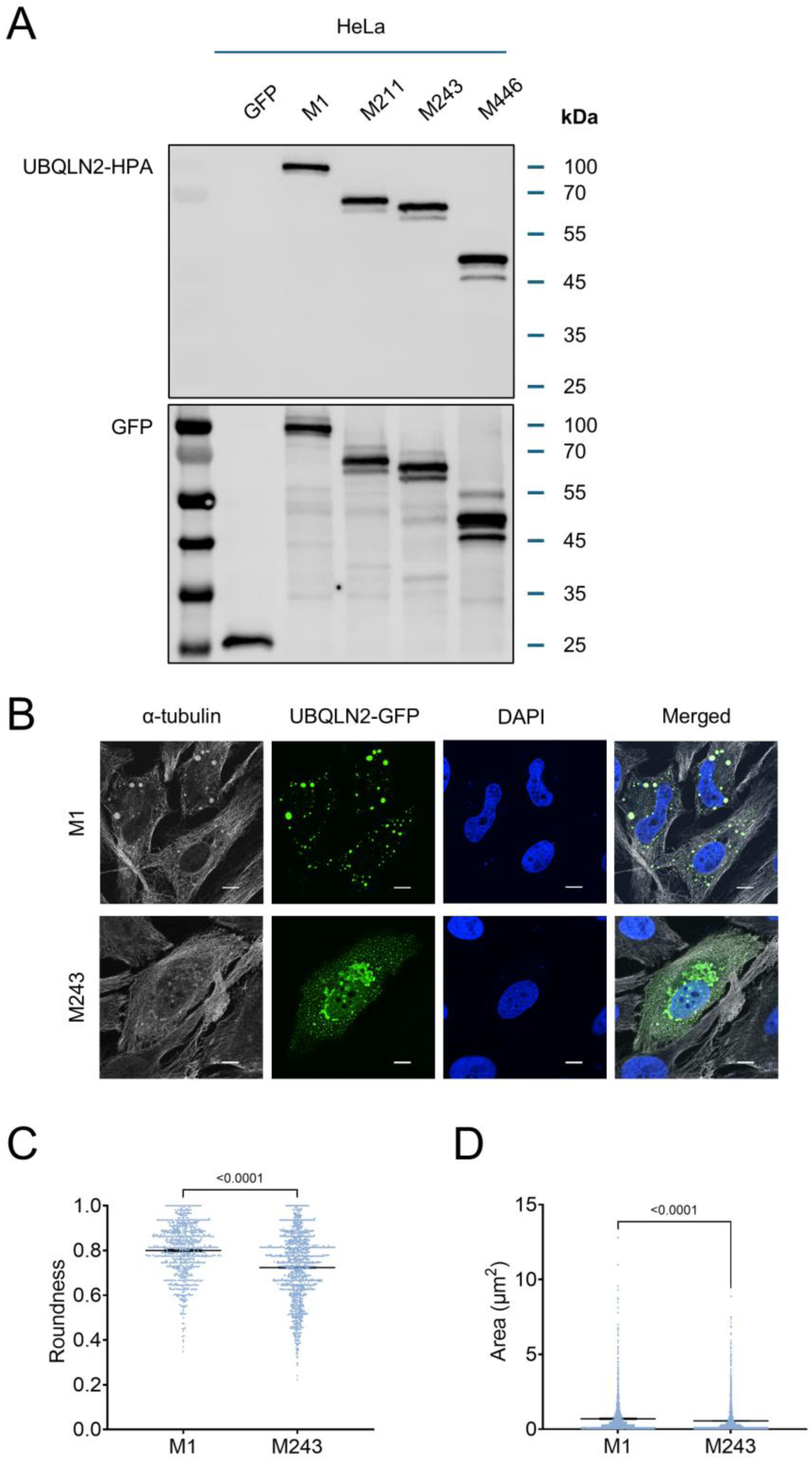
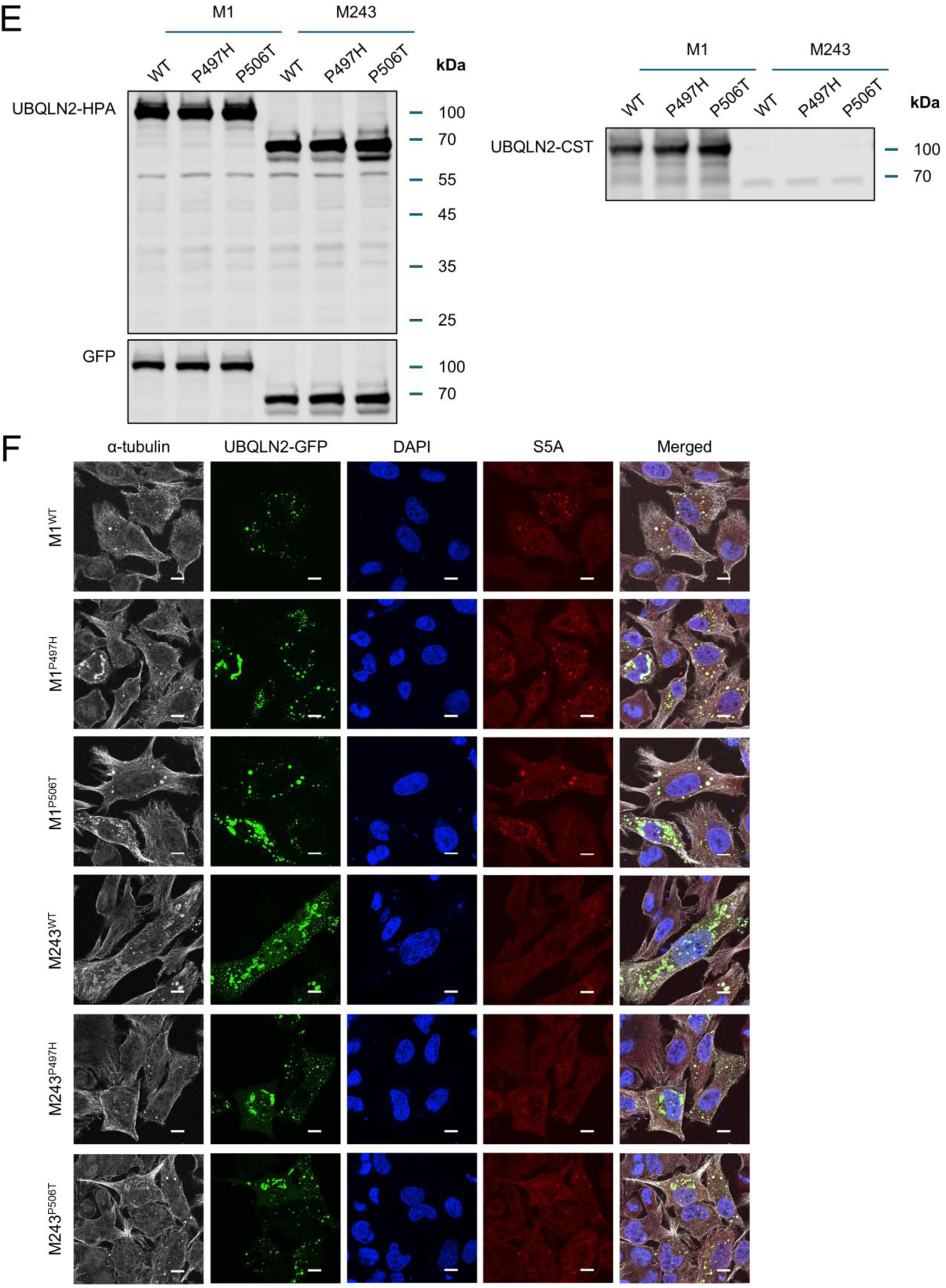
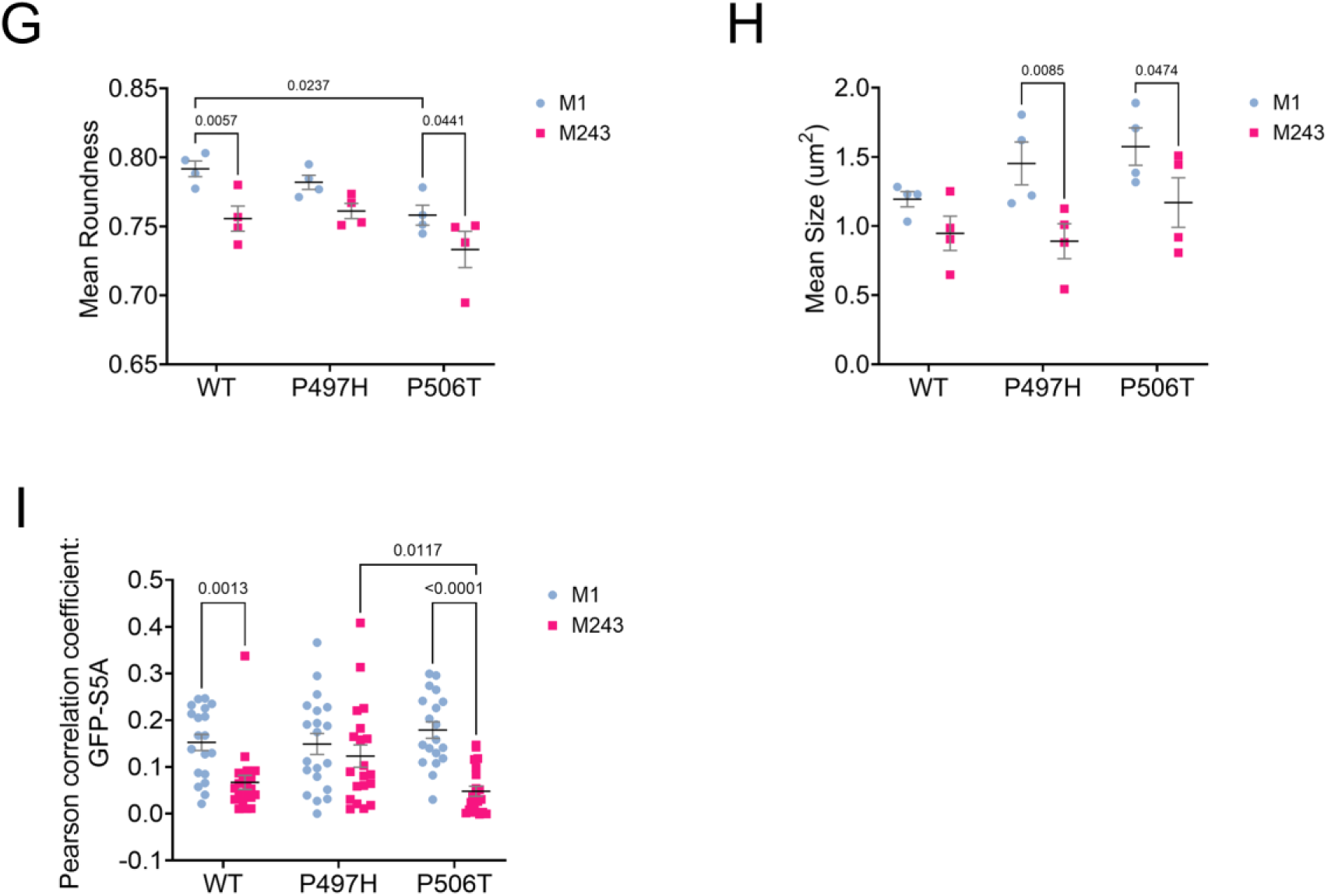
Relative expression and localization of GFP-UBQLN2 isoforms and their ALS-associated mutants. (A) Representative Western blot images of HeLa cells transiently transfected with pcDNA5-eGFP plasmids expressing the corresponding UBQLN2 isoforms. (B) HeLa cells expressing GFP-tagged UBQLN2-M1 and M243 isoforms were fixed and stained with α-tubulin. Cell nuclei were counterstained with DAPI. Scale bar = 10 µm. (C) Quantification of the roundness of the aggregates (larger than 0.2 µm^2^) formed by cells in (B). Full-length UBQLN2 (M1) forms spherical cytosolic aggregates compared to the irregular aggregates formed by M243 isoforms. Mean ± SEM were shown. Each dot represents measurement from an aggregate tabulated from one biological replicate, number of aggregates for M1 = 852 and M243 = 1205. *p*-value from two-tailed unpaired *t*-test was listed. The experiment was repeated four times in HeLa cells which reproduced similar observations; aggregate roundness quantification from three other replicates were shown in Sup. Fig. 4A-C. (D) Tabulation of the aggregate size from cells expressing indicated UBQLN2 constructs. Mean ± SEM were shown. Each dot represents measurement from an aggregate tabulated from one biological replicate, number of aggregates for M1 = 1429 and M243 = 2454. *p*-value from two-tailed unpaired *t*-test was listed. (E) Representative Western blot images of HeLa cells transiently transfected with pcDNA5-eGFP plasmids expressing the GFP-tagged UBQLN2-M1 and M243 isoforms with the indicated ALS mutations. (F) HeLa cells expressing indicated UBQLN2 isoforms and their corresponding ALS mutants were fixed and stained with α-tubulin and proteosome marker S5A. Cell nuclei were counterstained with DAPI. Scale bar = 10 µm. (G) Quantification of the mean roundness of the aggregates (larger than 0.2 µm^2^) formed by cells in (F). P506T mutants were more prone to irregular aggregate formation. Mean ± SEM were shown. Each dot represents mean aggregate roundness value from one biological replicate, N = 4. Plots for individual biological replicates were shown in Sup. Fig. 4D-G. Significant *p*-values from two-way ANOVA and Tukey’s multiple comparisons test were listed. (H) Tabulation of the mean aggregate size formed by cells in (F). Mean ± SEM were shown. Each dot represents mean aggregate size from one biological replicate, N = 4. Plots for individual biological replicates were shown in Sup. Fig. 4H-K. Significant *p*-values from two-way ANOVA and Tukey’s multiple comparisons test were listed. (I) Pearson correlation coefficient of the GFP and S5A masks was calculated for each image taken from a total of three biological replicates. Mean ± SEM were shown. Each dot represents an image, 19 ≤ N ≤ 21. Significant *p*-values from two-way ANOVA and Šidák’s multiple comparisons test were listed.

We next compared the expression and localization patterns of GFP-UBQLN2-M1 and GFP-UBQLN2-M243 proteins harboring ALS-associated P497H and P506T mutations (Fig. 6E,F). UBQLN2-M1^P506T^ and UBQLN2-M243^P506T^ formed aggregates with a significant reduction in roundness compared to the corresponding wild-type constructs (Fig. 6F,G, Sup. Fig. 4D-G). By contrast, P497H mutations on both UBQLN2-M1 and UBQLN2-M243 backgrounds did not have a significant effect on roundness. Both P497H and P506T mutations also significantly reduced the aggregate size of the UBQLN2-M243 isoform compared to the corresponding mutant in UBQLN2-M1 (Fig. 6F,H, Sup. Fig. 4H-K).

As UBQLN2-M243 lacks the UBL proteosome-targeting domain, we examined the colocalization of UBQLN2-M243 aggregates with the proteasome marker, S5A (Fig. 6F,I). Pearson correlation coefficient analysis demonstrated reduced colocalization of UBQLN2-M243^WT^ and UBQLN2-M243^P506T^ with S5A relative to their UBQLN2-MI counterparts (Fig. 6I). These findings suggest that the UBQLN2-M243 isoform produced from the spliced UBQLN2 transcripts has altered functional characteristics that may be differentially impacted by ALS-associated mutations.

### *UBQLN2* splicing is upregulated in male ALS patients

Altered exitron splicing has the potential to impact *UBQLN2* gene expression and, potentially, proteostasis, in ALS/FTD. To investigate this, we queried a subset of TargetALS transcriptomic data from postmortem medial motor cortices (Sup. Table 1) for an estimation of the UBQLN2-Sp isoform expression. This analysis revealed a significant increase in the mean UBQLN2-Sp isoform percentage from ∼6% in non-neurological controls to ∼10% in the medial motor cortex of male ALS patients, with some patient samples exhibiting ∼25-30% UBQLN2-Sp isoform percentage (Fig. 7A). UBQLN2-Sp isoform levels were also significantly higher in male ALS patients versus female ALS patients, but there was not significantly different between female ALS patients and the non-neurological controls (Fig. 7A). Higher UBQLN2-Sp isoform expression was weakly correlated to lower *UBQLN2* gene expression in males (Fig. 7B,C). Combined with the finding that TDP-43 knockdown increased *UBQLN2* splicing in cell culture (Fig. 2B,C), these results suggest that *UBQLN2* exitron splicing is increased during the clinical course of neurodegeneration in male ALS/FTD patients and that exitron splicing may be inversely correlated to *UBQLN2* gene dosage.

**Fig. 7.**
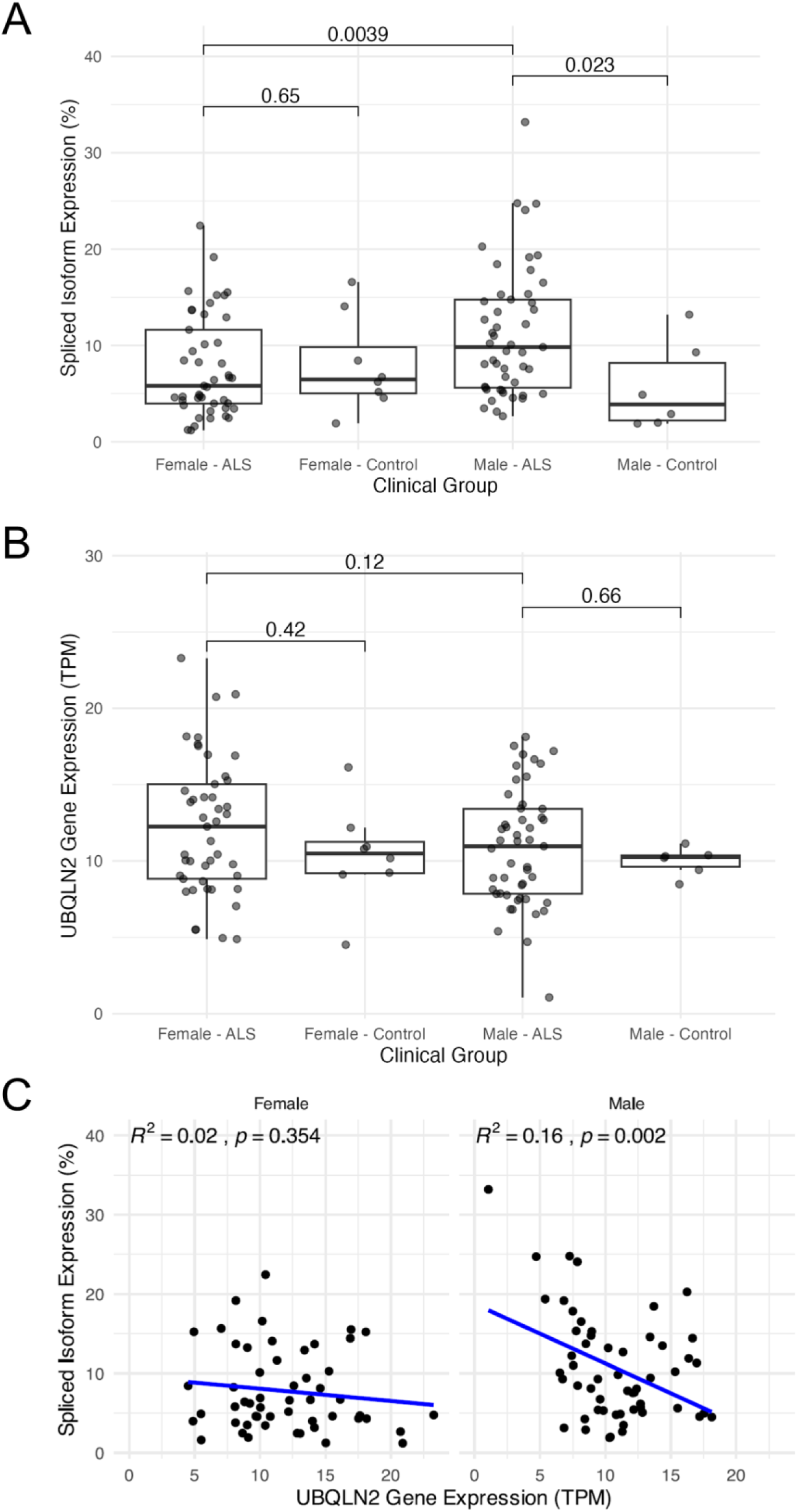
*UBQLN2* exitron splicing is elevated in ALS. (A) Transcriptomes of ALS patients (medial motor cortex, female = 45; male = 49) and non-neurological controls (female = 8; male = 6) were retrieved from the TargetALS database (see Sup. Table 1 for ExternalSampleIDs). UBQLN2-Spliced Isoform Expression (%) was estimated by RSEM. *p*-values from Wilcoxon test were listed. (B) *UBQLN2* gene expression (TPM) from ALS transcriptomes. *p*-values from Wilcoxon test were listed. (C) Correlation between UBQLN2-Spliced Isoform Expression (%) (A) and UBQLN2 gene expression (TPM) (B).

## DISCUSSION

UBQLN2-ALS pathogenesis likely involves multiple pathways related to both toxic gain of function (GOF) and loss of function (LOF) mechanisms ^101^. Furthermore, because both *UBQLN2* overexpression and silencing cause neurological phenotypes in rodents ^58,98,102^, perturbation of wild-type *UBQLN2* expression levels may have implications for neuronal proteostasis and neurodegeneration. Here we demonstrate that the expression of full-length *UBQLN2* is influenced by an unannotated exitron whose splice out produces an N-terminally truncated UBQLN2-M243 isoform lacking the UBL domain which is unstable and showed distinct aggregation phenotypes when overexpressed. *HsUBQLN2* exitron splicing is inhibited by heat shock (Fig. 2I,J) and in differentiating neurons (Fig. 2G,H) which may have a strong requirement for UBQLN2-mediated proteostasis. *UBQLN2* splicing was also elevated in ALS medial motor cortex in males (Fig. 7). These findings suggest the possibility that alternative splicing regulates functional *HsUBQLN2* gene dosage and that truncated UBQLN2 isoforms may uniquely contribute to the neuropathogenesis of ALS/FTD-UBQLN2.

Exitrons were first discovered in *Arabidopsis* and can arise from the activation of cryptic splice sites within protein-coding exons ^92,103,104^. While originally estimated to affect ∼5-10% of coding genes, recent studies suggest that exitron splicing is more extensive than previously thought, occurring in ∼63% of genes from cancer-derived cell lines ^105^. Individual exitron splicing events showed strong tumor specificity and may be related to global RBP dysregulation in cancer ^105^. In gastric cancers driven by ALS-related splicing factors such as FUS and TAF15, *UBQLN2* was one of the more than 1,500 significantly spliced genes detected ^106^. Exitron regulation in ALS/FTD has not been explored; however, given the centrality of RNA splicing defects in these diseases, it is possible that alterations in exitron removal contribute to gene expression changes, protein aggregation, and neurodegeneration.

Using minigene reporters, we identified a single SD and two SAs that mediate *HsUBQLN2* exitron splicing. SA2 is the major SA used in human cells, but SA1 and other upstream unidentified SAs could be used when SA2 was mutated (Fig. 4B-D). Interestingly, despite the fact that both SD and SA sequences are conserved between *MmUBQLN2* and *HsUBQLN2*, the PSO of UBQLN2 exitron was ∼100-fold higher in human cells versus mouse cells (Fig. 1) and ∼10-50-fold higher for the *HsUBQLN2* reporter versus the *MmUBQLN2* reporter in HeLa cells (Fig. 3B-F). suggesting that exitron splicing arose after the evolutionary divergence of primates and mice from their common ancestor.

Experiments using chimeric minigenes revealed that sequence elements in the 5’UTR contribute to the different splicing efficiencies of *HsUBQLN2* and *MmUBQLN2* in human cells. Knockdown, mutagenesis, overexpression, and RNA-IP studies further suggest that PTBP1 (and potentially PTBP2 in differentiating neurons) can both activate and repress *HsUBQLN2* exitron splicing through binding to Py motifs located in the 5’UTR (Fig. 2A-F, Fig. 4). PTBPs fulfill complex roles in alternative splicing depending on splice site usage frequency and positional context. In general, binding of PTBPs near a constitutive/strong splice site promotes exon inclusion whereas binding near an alternative splice site often causes exon skipping ^40^. While PTBP1 often binds to intronic region upstream of SAs, binding to sequence downstream of an SD has been observed, such as PTBP2-Exon 10, whose splicing is repressed by PTBP1 ^40^. The human-specific Py2 (C<u>UCUCUCU</u>UUC) contains a known binding consensus which is shared amongst PTBP paralogs: UCUC/UUCU (Fig. 2A). Consistent with our PTBP1 RNA-IP in HeLa cells, Dawicki-McKenna *et al.* detected a PTBP2 CLIP-seq peak (chrX:56563827-56563877) in human iPSC-derived neurons ^41^, overlapping Py2 and Py3 tracts. Deletion of Py2 enhanced *HsUBQLN2* splicing efficiency (Fig. 4A-C), supporting its role as a major PTBP binding site that suppresses exitron removal. Remarkably, deletion of the adjacent Py1 motif, which contains a tandemly duplicated CCTCCTTCC motif in *HsUBQLN2* versus a single CCTCCTTCC motif in *MmUBQLN2*, reduced exitron splicing (Fig. 4A,G,H), as did a Py1/2 double mutant. Deletion of Py3, which is conserved between *HsUBQLN2* and *MmUBQLN2*, also diminished splicing, suggesting this element may be important for the low levels of MmUBQLN2 splicing that we observed (Fig. 3A-F).

The fact that Py motifs upstream and downstream of the SD in the 5’UTR have opposing impacts on *UBQLN2* splicing raises the question as to how these sites interact to control *UBQLN2* splicing efficiency. One possibility is that the overall impact of PTBPs on *UBQLN2* exitron splicing is determined by combinatorial effects of Py tracts that are occupied with distinct stoichiometries and, perhaps under distinct cellular conditions. For instance, low levels of PTBP1 may result in selective occupancy of high affinity sites (putatively Py2) to inhibit SD recognition, while high levels of PTBP1 during overexpression (Fig. 4K,L) may promote occupancy of both high affinity and low affinity (putatively Py1 and Py3) sites which enhance splicing. In the latter case, the established multimerization activities of PTBP1 ^38^ may alter the secondary structure of *HsUBQLN2* 5’UTR. Alternatively, multiple engagement of its four RNA recognition motifs which are structurally heterogeneous ^107^ with different affinities to Py2 and Py1/3 may control the stoichiometries of bound PTBP1 to influence splicing outcome. Future studies will be needed to define structural features of the *UBQLN2* 5’UTR and its combinatorial regulation by PTBPs.

The implication of SRSF1 as a *UBQLN2* splicing repressor (Fig. 2A-E; Fig. 4K,L) which bound to Pu2 (Fig. 4A,I,J) is supported by the detection of an SRSF1 eCLIP peak in the 5’UTR of UBQLN2 in a previous study in K562 cells ^108^. Mutations of another downstream putative SRSF1 binding site, Pu1, inhibited *UBQLN2* splicing (Fig. 4A,I,J), suggesting this element function as a splicing enhancer and could be occupied by either SRSF1 or other splicing enhancers. Finally, we also observed that PTBP1 expression level is inversely correlated with SRSF1 expression (Fig. 2D, 4M), suggesting that *HsUBQLN2* exitron splicing is regulated through complex interplay of PTBP1, SRSF1, and other RBPs, with TDP-43 being of particular interest. We note that in HeLa cells, the expression of TDP-43 stays relatively constant, therefore, further studies are needed to investigate the contributions of PTBP1, PTBP2, SRSF1, and TDP-43 to *UBQLN2* splicing regulation in neurons.

Spliced *HsUBQLN2* transcripts express UBQLN2-M243 and potentially other N-terminally truncated isoforms (Fig. 5A-D,G). Consistent with the absence of a UBL domain, UBQLN2-M243 showed reduced colocalization with proteasome markers (Fig. 6F,I), suggesting its function in UPS is substantially altered relative to full-length UBQLN2. UBQLN2 STI1 domains are the main driver for its LLPS propensity, and the removal of the UBL domain has been shown to reduce the saturation concentration for UBQLN2 LLPS ^53,109^. Indeed, UBQLN2-M243 formed smaller and more irregular aggregates versus UBQLN2-M1, suggesting altered phase separation characteristics (Fig. 6, Sup. Fig. 4). Amongst all ALS mutations, UBQLN2^P506T^ was found to have the highest propensity in intrinsic self-assembly to form amyloid-like aggregate through LLPS ^110,111^. *UBQLN2* exitron splicing can potentially exacerbate UBQLN2^P506T^ LLPS characteristics in UBQLN2-M243 compared to UBQLN2-M1 (Fig. 6F-H). Although comprising less than 2% of total UBQLN2 expressed from the *HsUBQLN2* minigene at steady state (Fig. 5B,C) due to rapid degradation by proteasome (Fig. 5E,F), the altered phase separation and function of UBQLN2-M243 may be relevant to the regulation of cellular proteostasis and the neuropathogenesis of ALS/FTD-UBQLN2, especially in the case where protein clearance pathways are impaired in NDs.

While the contribution of UBQLN2 isoforms to ALS/FTD-UBQLN2 and sporadic ALS is still uncertain, we observed elevated levels of splicing in motor cortices of male ALS patients (Fig. 7). This is interesting given the X-linkage of *UBQLN2* and a recent metaanalysis showing that males harboring a pathogenic *UBQLN2* variant developed ALS on average 18.15 years prior to female carriers (29.54 ± 11.9 versus 47.69 ± 13.4 years, p < 0.0001) ^112^. In our analysis, increased *UBQLN2* splicing in male ALS patients was also associated with reduced *UBQLN2* gene expression, raising the possibility that *UBQLN2* exitron splicing dysregulation contributes to proteostasis failure through LOF effects. A consequence of proteostasis failure—the accumulation of cytosolic TDP-43—may further exacerbate *UBQLN2* splicing dysregulation, leading to feed-forward toxicities that drive disease progression in ALS/FTD.

## ACKNOWLEDGEMENTS

We thank Dr. Ong Swee Hoe (Wellcome Sanger Institute) for suggestions to improve our RNA sequencing analysis pipeline, Dr. Stephen Turner (Pacific Biosciences) and Dr. Paola Florez de Sessions (Oxford Nanopore Technologies) for assistance in sourcing publicly available long-read RNA sequencing datasets. We acknowledge Jeremy Niece from UWBC for speedy processing of our sequencing samples.

## AUTHOR CONTRIBUTIONS

Conceptualization, R.S.T., A.S.H.K., J.Z.;

Methodology, all;

Investigation, Validation, and Formal Analysis, A.S.H.K., J.Z., N.M-A., Y.W.;

Software, Data Curation, and Visualization, A.S.H.K., J.Z., N.M-A.;

Writing – Original Draft, R.S.T., A.S.H.K.;

Writing – Review & Editing, all;

Supervision, Resources, and Funding Acquisition, R.S.T., X.Z., W.X.;

## DATA AVAILABILITY

The data underlying this article will be shared on reasonable request to the corresponding author.

