## Supplemental Figures for "ALS-associated exitron splicing produces UBQLN2 isoforms with distinct properties"

### SUPPLEMENTARY FIGURES

Koo *et al.* Sup. Fig. 1

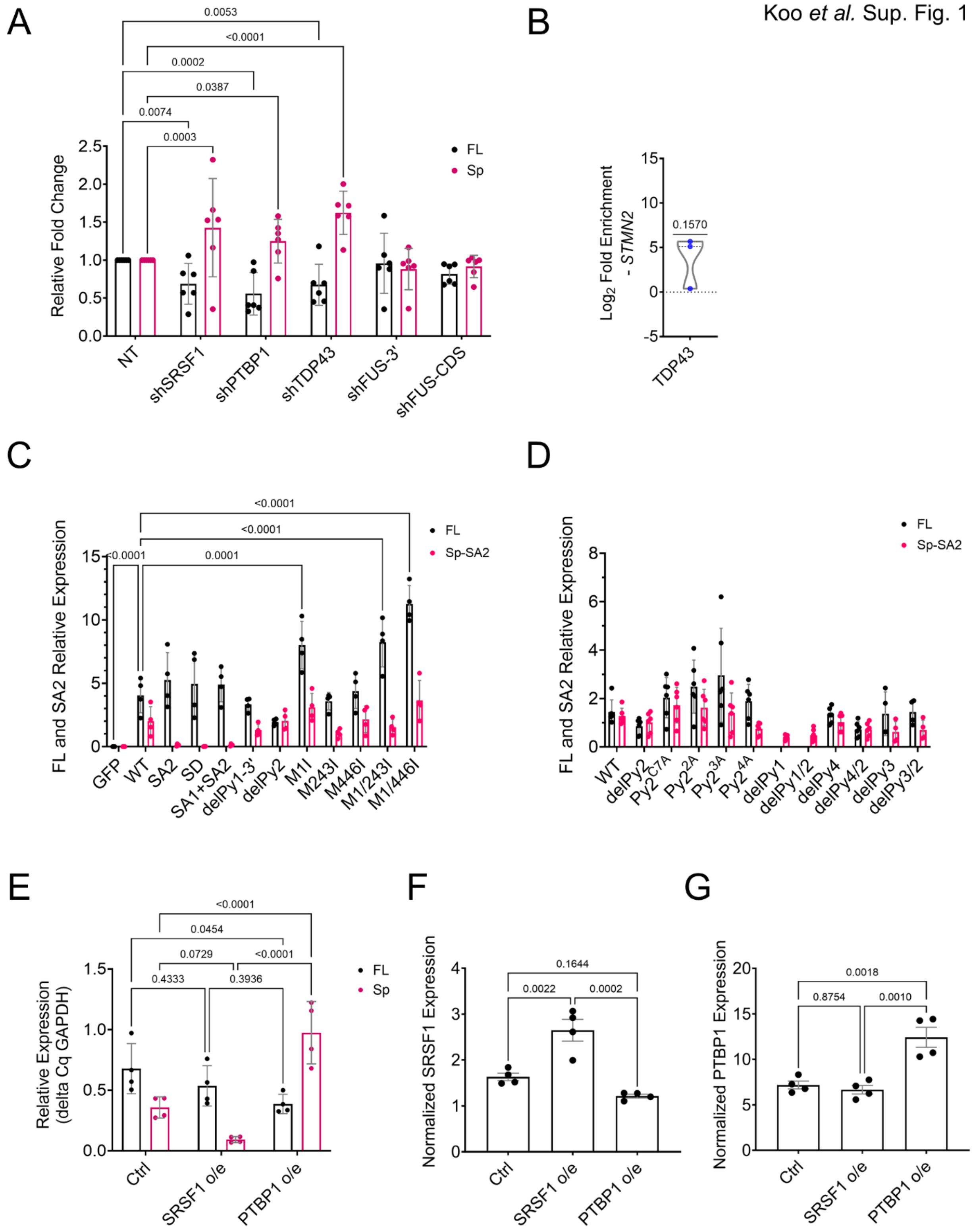

**Sup. Fig. 1. *UBQLN2* splicing is a result of both direct and indirect regulation by PTBP1 and SRSF1.**

(A) Quantification of the relative fold change (ddCq) of full-length UBQLN2-FL and UBQLN2-Sp in HeLa cells transduced with shRNA vectors targeting the indicated RBPs by qRT-PCR. The Cq values of the FL and Sp were normalized to GAPDH and its corresponding baseline Cq value of non-targeting control (NT). Bar height corresponds to mean fold change ± SEM. Each dot represents a biological replicate, N = 6, significant *p*-values from repeated measures two-way ANOVA and Dunnett's multiple comparisons test were listed.

(B) $Log_2$ fold enrichment of STMN2 pre-mRNA immunoprecipitated by TDP-43 antibody in native RNA immunoprecipitation assays in HeLa cells. Median and interquartile range were shown as dotted lines. Each dot represents an individual biological replicate, N = 3. *p*-value from two-tailed one sample t-test was listed.

(C) Quantification of the relative expression of UBQLN2-FL and UBQLN2-Sp (SA2) transcripts for the PSO calculation in Fig. 4C. Significant *p*-values from two-way ANOVA and Dunnett's multiple comparisons test were listed.

(D) Quantification of the relative expression of UBQLN2-FL and UBQLN2-Sp (SA2) transcripts for the PSO calculation in Fig. 4H. UBQLN2-FL transcripts could not be amplified in delPy1 and delPy1/2 due to a deletion which affects qPCR primer binding.

(E) Quantification of the relative expression of UBQLN2-FL and UBQLN2-Sp in HeLa cells overexpressing SRSF1 or PTBP1 as shown in Fig. 4I-J. The Cq values of the FL and Sp were normalized to GAPDH. Bar height corresponds to mean ± SEM. Each dot represents a biological replicate, N = 4, significant *p*-values from two-way ANOVA and Tukey's multiple comparisons test were listed.

(F, G) SRSF1 and PTBP1 expression were quantified based on densitometry from Fig. 4K and normalized to MCM2 or vinculin. Note that PTBP1 o/e caused a slight decrease in SRSF1 expression. Each dot represents a biological replicate, N = 4, significant *p*-values from one-way ANOVA and Tukey's multiple comparisons test were listed.

A

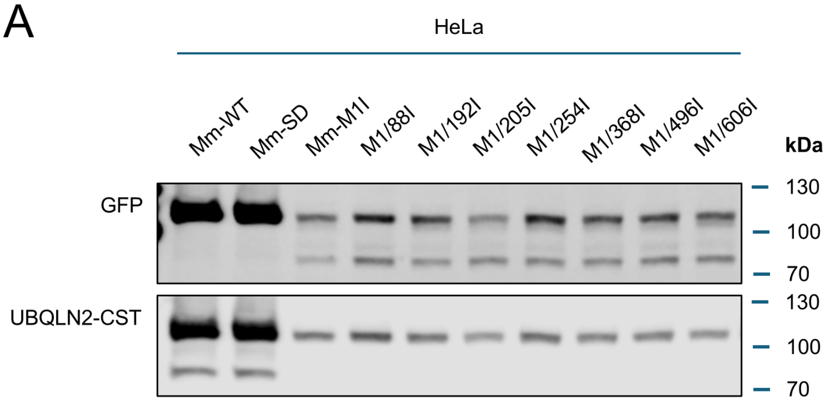


**Sup. Fig. 2. Mutation of candidate AUG codons failed to identify a major TSS downstream of M1 in MmUBQLN2.**

(A) AUG to AUC substitutions at codons 88, 192, 205, 254, 368, 496, and 606 were introduced into the MmUBQLN2-M1I splicing reporter that can only initiate translation downstream of M1. None of these mutations abolished the 75-kDa species that were observed in UBQLN2-M1I mutant cells.

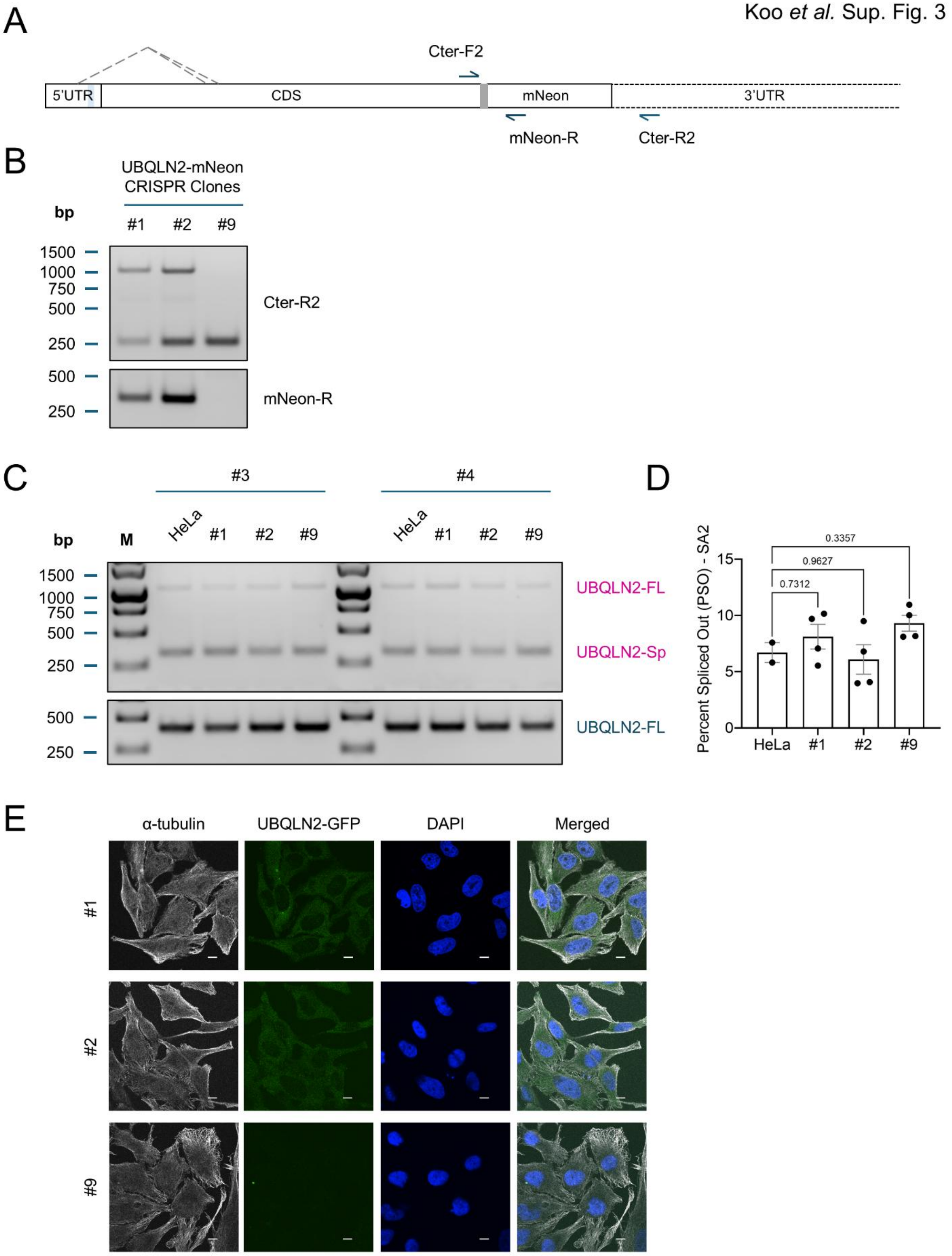


**Sup. Fig. 3. Characterization of UBQLN2-mNeon knockin HeLa cells.**

(A) Schematic showing mNeon knock-in at the C-terminus of UBQLN2 in HeLa cells. *Gray box* represents the 3x GGGGS linker inserted between UBQLN2 and mNeon CDS. The relative positions of the genotyping primers were indicated. Due to the length of the 3'UTR, it was not shown completely in the schematic and was presented with *dotted lines*.
(B) Genotyping result of clones #1 and #2 which are positive for mNeon insertion. Clone #1 and #2 contain an mNeon insertion in at least one allele. Clone #9 was included as a negative control.
(C, D) *UBQLN2* splicing assay showed that CRISPR generated clones have comparable UBQLN2 exitron PSO as the unedited HeLa cells. Bar height corresponds to mean ± SEM. Each dot represents a biological replicate, N = 4 for each CRISPR clones, N = 2 for unedited HeLa cells. *p*-values from one-way ANOVA and Dunnett's multiple comparisons test were listed.
(E) Confocal images showing UBQLN2-mNeon expression in clones #1 and #2. Cells were fixed and stained with α-tubulin. Cell nuclei were counterstained with DAPI. Scale bar = 10 μm.

Koo *et al.* Sup. Fig. 4

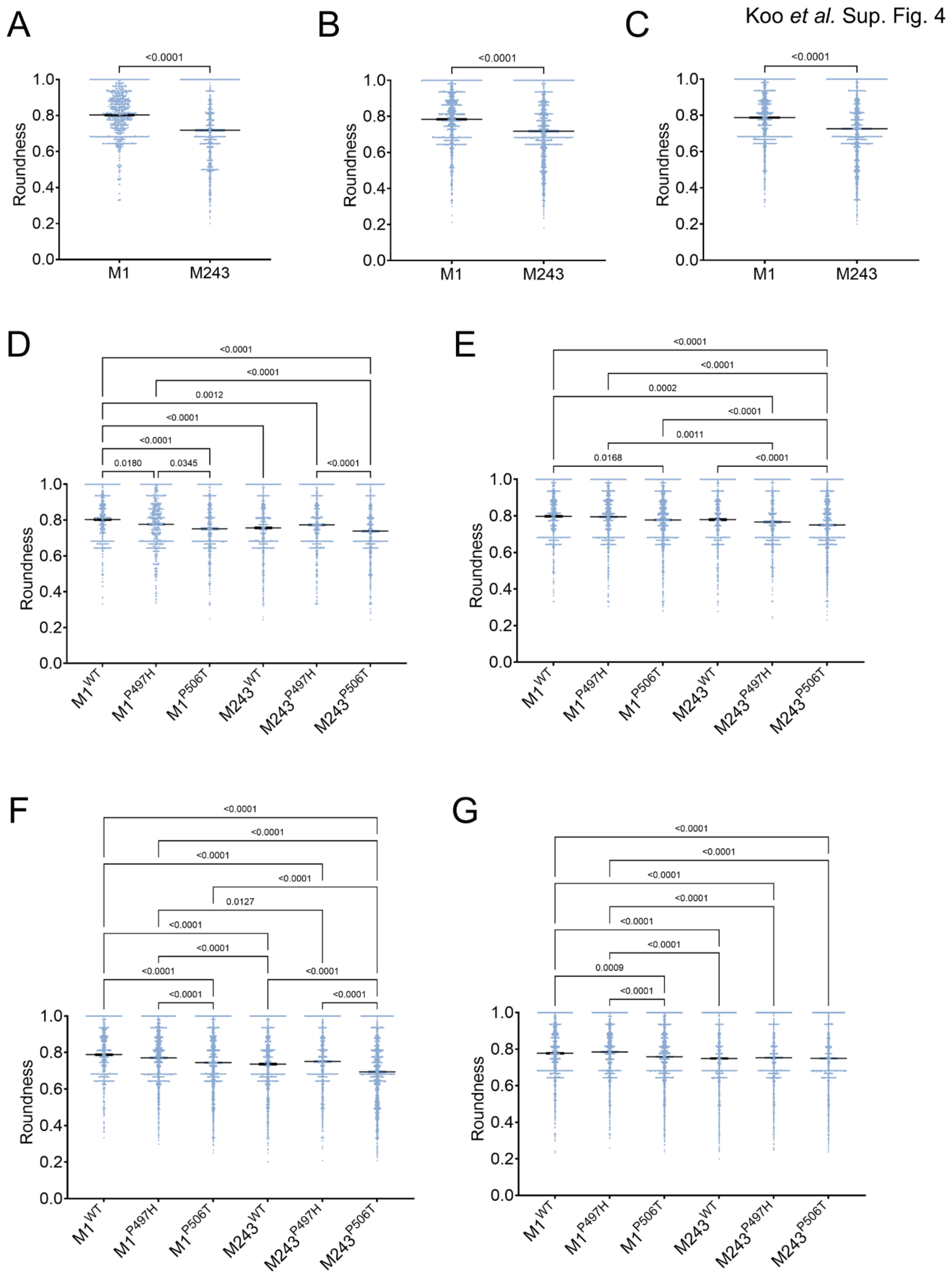

H

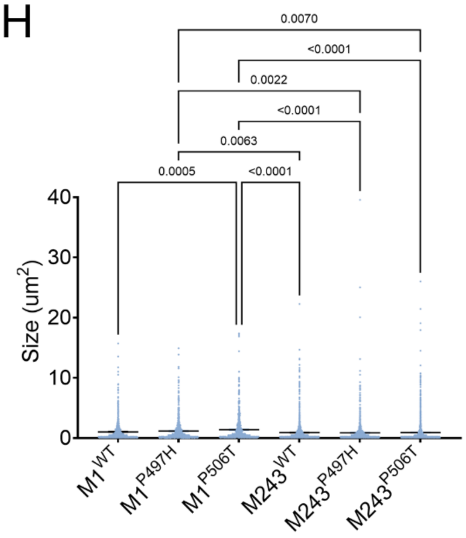


I

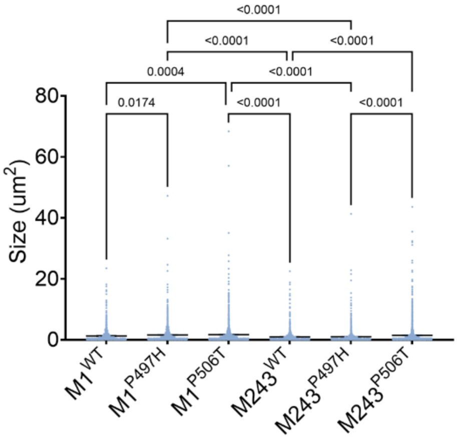


J

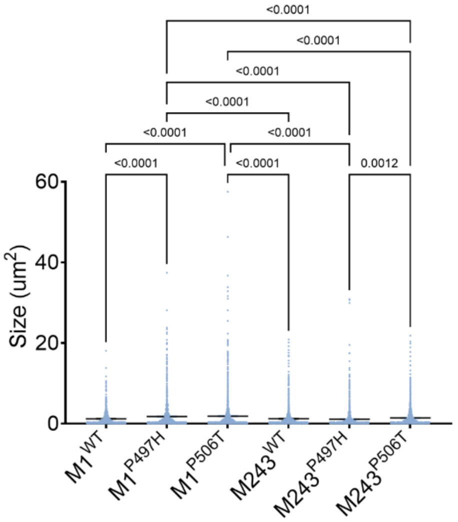


K

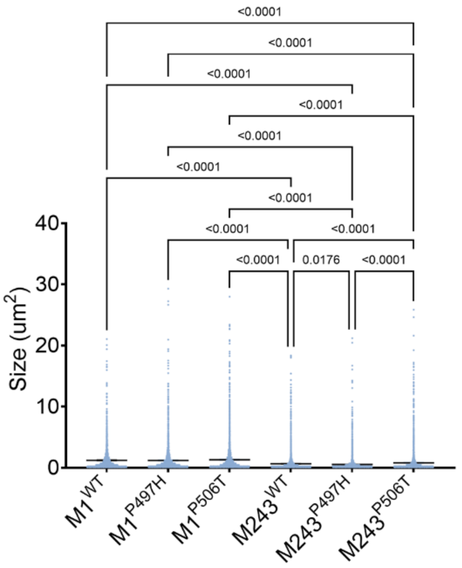


**Sup. Fig. 4. Quantification of aggregate roundness and size of UBQLN2 isoforms and their ALS-associated mutants.**

(A-C) Quantification of the aggregate roundness (larger than 0.2 μm$^2$) formed by cells in experimental replicates #2 - #4 for Fig. 6C. Replicate #2: number of aggregates for M1 = 1003 and M243 = 1560; Replicate #3: number of aggregates for M1 = 1540 and M243 = 1619; Replicate #4: number of aggregates for M1 = 2076 and M243 = 3122. *p*-value from two-tailed unpaired *t*-test was listed.

(D-G) Quantification of the aggregate roundness (larger than 0.2 μm$^2$) formed by cells in experimental replicates #1 - #4 for Fig. 6F. Replicate #1: 608 ≤ number of aggregates ≤ 1191; Replicate #2: 784 ≤ number of aggregates ≤ 1583; Replicate #3: 978 ≤ number of aggregates ≤ 2031; Replicate #4: 1785 ≤ number of aggregates ≤ 3081. Significant *p*-values from one-way ANOVA and Tukey's multiple comparisons test were listed.

(H-K) Quantification of the aggregate size formed by cells in experimental replicates #1 - #4 for Fig. 6G. Replicate #1: 780 ≤ number of aggregates ≤ 2057; Replicate #2: 1139 ≤ number of aggregates ≤ 2132; Replicate #3: 1265 ≤ number of aggregates ≤ 2550; Replicate #4: 2382 ≤ number of aggregates ≤ 5518. Significant *p*-values from one-way ANOVA and Tukey's multiple comparisons test were listed.
